# Ion channel function of polycystin-2/polycystin-1 heteromer revealed by structure-guided mutagenesis

**DOI:** 10.1101/2024.11.22.624889

**Authors:** Tobias Staudner, Juthamas Khamseekaew, M. Gregor Madej, Linda Geiges, Bardha Azemi, Christine Ziegler, Christoph Korbmacher, Alexandr V. Ilyaskin

## Abstract

Autosomal-dominant polycystic kidney disease (ADPKD) is caused by mutations affecting polycystin-1 (PC1) or polycystin-2 (PC2). Recent structural data suggest that PC1 and PC2 can form heterotetrameric ion channels with a 3:1 stoichiometry, with the channel in a closed state. In this hetero-oligomeric formation three PC1 residues (R4100, R4107, and H4111) would work together with two PC2 residues (L677, N681) to block the ion permeation pathway. Here, we provide functional evidence supporting this model. When the pore-blocking residues are replaced by alanines, there is a significant gain-of-function (GOF) effect on PC2/PC1 currents. These PC2/PC1 GOF channels show different selectivity for monovalent cations and reduced Ca^2+^ permeability compared to PC2 homomers. By varying the expression ratio of PC1 and PC2, we found that PC2 preferentially forms heteromeric complexes with PC1 over homomers. Additionally, a re-interpretation of published PC2/PC1 cryo-electron microscopy data, combined with cysteine modification experiments, suggests that the pore-forming domain of PC1 adopts a canonical TRP-like conformation. The novel PC2/PC1 GOF construct offers a promising approach to investigate the functional impact of mutations linked to ADPKD.

## Introduction

Autosomal dominant polycystic kidney disease (ADPKD) is the most common hereditary kidney disease and is responsible for 5-10% of all cases of end-stage renal disease (ESRD) (Chebib & Torres, 2016). Its main manifestation is the development and progressive growth of multiple fluid-filled renal cysts which compromise kidney function (Grantham *et al*, 2011). In the majority of cases, ADPKD is caused by mutations in the *PKD1* or *PKD2* gene (Bergmann *et al*, 2018; Cornec-Le Gall *et al*, 2018) coding for polycystin-1 (PC1) and polycystin-2 (PC2), respectively (Hughes *et al*, 1995; Mochizuki *et al*, 1996). On a cellular level ADPKD-onset is most likely triggered by a two-hit mechanism requiring one germline mutation (“first hit”) and a second somatic mutation (“second hit”) (Koptides *et al*, 1999; Pei *et al*, 1999; Qian *et al*, 1996; Tan *et al*, 2018; Wu *et al*, 1998). Despite intensive research efforts, the physiological function of PC1 and PC2 and their mechanistic role in ADPKD pathogenesis remain unclear.

PC2 is a member of the transient receptor potential (TRP) family of ion channels (Wu *et al*, 2010) bearing its hallmark features of 6 transmembrane domains (S1-S6), intracellular N-and C-termini, and a putative ion permeation pathway between the fifth (S5) and the sixth (S6) transmembrane domain (Hoffmeister *et al*, 2011). Recently published cryo-electron microscopy (cryo-EM) structures of PC2 indicate that the protein can form homotetrameric ion channels (Grieben *et al*, 2017; Shen *et al*, 2016; Wang *et al*, 2020a; Wang *et al*, 2024; Wilkes *et al*, 2017). Therefore, homotetrameric PC2 probably functions as non-selective cation channel like most TRP channels (Cao, 2020; Wu *et al*, 2010). However, both the (patho-)physiological role and the electrophysiological characteristics of PC2 remain controversial in the literature (Cantero & Cantiello, 2022; Douguet *et al*, 2019). Early studies identified the endoplasmic reticulum as one of the main sites of PC2 expression (Cai *et al*, 1999; Koulen *et al*, 2002). Therefore, it was proposed that PC2 either mediates Ca^2+^ release from the endoplasmic reticulum (Koulen *et al*., 2002; Wegierski *et al*, 2009) or interacts with ryanodine receptors (Anyatonwu *et al*, 2007) and IP_3_-receptors (Li *et al*, 2005; Sammels *et al*, 2010). A more recent study proposed that PC2 might function as a K^+^ ion channel in the ER facilitating K^+^-Ca^2+^ counterion exchange, thereby promoting IP_3_-mediated Ca^2+^ release from ER (Padhy *et al*, 2022). Another cellular location of PC2 is the primary cilium (Pazour *et al*, 2002; Yoder *et al*, 2002). Primary cilia malfunction is thought to contribute to ADPKD pathogenesis (Ma *et al*, 2017; Ma *et al*, 2013; Walker *et al*, 2019), and a ciliary ion channel conductance was attributed to PC2 (Kleene & Kleene, 2017; Liu *et al*, 2018). Due to the absence of specific activators and inhibitors of PC2, its functional characterization remains challenging. Furthermore, in heterologous expression systems PC2 produces very low baseline currents (Arif Pavel *et al*, 2016; Grosch *et al*, 2021; Staudner *et al*, 2024). In recent years, several gain-of-function (GOF) mutations of PC2 have been generated (Arif Pavel *et al*., 2016; Wang *et al*, 2019; Zheng *et al*, 2018). This facilitated the electrophysiological characterization of PC2 in heterologous expression systems and made it possible to study effects of ADPKD-associated mutations on PC2 ion channel function (Staudner *et al*, 2024; Wang *et al*, 2023).

PC1 is a 4,303 residue-long transmembrane protein consisting of 11 transmembrane segments, a large ∼3,000 residue-long extracellular N-terminus and a short intracellular C-terminus (Hughes *et al*., 1995; Nims *et al*, 2003). The N-terminus of PC1 undergoes autocatalytic cleavage between L3048 and T3049 at the G protein-coupled receptor (GPCR) proteolysis site (GPS) of its GPCR autoproteolysis-inducing (GAIN) domain (Qian *et al*, 2002; Wei *et al*, 2007). This produces the N-terminal (PC1-NTF) and the C-terminal fragment (PC1-CTF), which remain non-covalently bound to each other. Because of these properties, PC1 resembles adhesion GPCRs (Bjarnadóttir *et al*, 2007; Nieberler *et al*, 2016). Furthermore, a putative tethered peptide agonist (“Stachel” domain) in the N-terminal portion of PC1-CTF (Pawnikar *et al*, 2024; Pawnikar *et al*, 2022) and a G-protein activation sequence in the intracellular C-terminus of PC1 have been described (Parnell *et al*, 2018; Parnell *et al*, 1998; Parnell *et al*, 2002). According to the generally accepted paradigm, PC1 and PC2 functionally interact *via* their C-terminal coiled-coil domains (Qian *et al*, 1997; Tsiokas *et al*, 1997; Yu *et al*, 2009; Zhu *et al*, 2011) and are co-localized in primary cilia (Yoder *et al*., 2002). This localization suggests an important yet unknown chemo-or mechanosensory function of the PC2/PC1 complex (Bergmann *et al*., 2018). However, it was demonstrated that PC1 is not required for basal ion channel activity of PC2 in primary cilia (Liu *et al*., 2018). Moreover, conflicting findings regarding the effect of PC1 on PC2 ion channel function in heterologous expression systems have been reported (Ha *et al*, 2024; Ha *et al*, 2020; Hanaoka *et al*, 2000; Wang *et al*., 2019). Therefore, it is still a matter of debate whether PC2/PC1 complex functions as an ion channel under physiological conditions and whether its disturbed ion channel function contributes to the onset and progression of ADPKD.

Recent cryo-EM data intriguingly indicate that PC1 can replace one PC2 subunit to form a heteromeric PC2/PC1 ion channel with a 3:1 stoichiometry (Su *et al*, 2018a). Moreover, co-expression of PC1 has been shown to alter ion channel properties of a gain-of-function PC2 construct suggesting that PC1 contributes to the channel pore (Wang *et al*., 2019). This finding challenged the classical view regarding the functional interaction of PC2 with PC1 (Caplan, 2019). In this study, we used structure-guided mutagenesis to provide further evidence for PC2/PC1 function as heteromeric ion channel. We generated a novel GOF PC2/PC1 ion channel and characterized its properties. Furthermore, we demonstrated that PC2 preferentially associates with PC1 to form heteromeric complexes, likely with a 3:1 stoichiometry consistent with the cryo-EM data. The initial interpretation of the cryo-EM densities suggested that PC1 has a noncanonical architecture in its pore-forming domains, where the S6 helix consists of two separate segments, S6a and S6b, with the proximal segment S6a resembling the pore helix 1 (Su *et al*., 2018a). However, our re-interpretation of these cryo-EM data did not confirm this structural feature. Instead, our revised model suggests that the pore-forming domain of PC1 has a canonical TRP-like conformation. Combined with our functional data, this supports the conclusion that PC1 directly contributes to the channel pore of the PC2/PC1 heteromeric ion channel. Moreover, the GOF PC2/PC1 construct developed in this research may serve as a valuable tool for elucidating PC2/PC1 ion channel properties and investigating the functional consequences of ADPKD-associated mutations in PC2 and PC1.

## Results and discussion

### Wild-type PC2 expressed alone or in combination with the C-terminal fragment of PC1 (PC1-CTF) did not reveal substantial ion channel activity in the oocyte expression system

To investigate the functional interaction between human PC2 and human PC1 we used *Xenopus laevis* oocytes as a heterologous expression system. Removal of the divalent cations (Ca^2+^ and Mg^2+^) from the bath solution was used as an established maneuver to reveal PC2 ion channel activity (Arif Pavel *et al*., 2016; Grosch *et al*., 2021; Staudner *et al*., 2024; Zheng *et al*., 2018). In oocytes expressing PC2 WT only marginal sodium inward currents could be detected under divalent cation free conditions (**PC2**; Fig. 1EV-A, D). These inward currents were completely blocked when extracellular Na^+^ was replaced by the large organic cation NMDG^+^ to which PC2 is essentially impermeable. These experiments demonstrated that the Na^+^ conductance of oocytes expressing PC2 WT was very small, consistent with our previous findings (Grosch *et al*., 2021; Staudner *et al*., 2024). It has been reported that the C-terminal fragment of PC1 (PC1-CTF) is sufficient to study PC2/PC1 interaction in the oocyte expression system (Wang *et al*., 2019). Co-expression of PC2 with PC1-CTF (amino acid residues 3049-4303; further referred to as **PC1**) reduced inward currents elicited by Ca^2+^ and Mg^2+^ removal (**PC2 + PC1**; Fig. 1EV-B and D) to a level below that in oocytes expressing PC2 alone and only slightly higher than that in control oocytes (Fig. 1EV-C, D). Importantly, decreased currents in this group of oocytes were not due to reduced PC2 expression at the cell surface. On the contrary, PC2 cell surface expression was increased in oocytes co-expressing PC2 and PC1 compared to oocytes expressing PC2 alone (Fig. 1EV-E; Appendix Fig. S1). This latter finding is consistent with a previously described stimulatory effect of PC1 on PC2 cell surface expression (Yu *et al*., 2009; Zhu *et al*., 2011). In summary, co-expression of PC1 reduced PC2 currents in oocytes. In contrast, an earlier study in CHO cells reported that cells co-expressing PC1 and PC2 showed higher currents than cells transfected with PC1 or PC2 alone (Hanaoka *et al*., 2000).

### Alanine substitutions of the pore blocking residues in PC2 and PC1 produced a novel gain-of-function (GOF) PC2/PC1 construct

Our functional results suggested that PC2 without or with PC1 co-expression stayed mainly in the closed state at the oocyte surface. Structural information indicates that in the closed state the ion permeation pathway of the PC2 homomeric channel is occluded by two lower gate residues L677 and N681 (Grieben *et al*., 2017; Shen *et al*., 2016; Wang *et al*, 2020b; Wilkes *et al*., 2017) Fig.1A). Similarly, in the heterotetrameric PC2/PC1 complex the putative lower gate is occluded by the same PC2 residues and additionally by three positively charged PC1 residues (R4100, R4107, H4111) (Su *et al*., 2018a), Fig.1B). It is now well-established that alanine substitutions of L677 and N681 in PC2 produce a strong GOF effect on PC2-mediated currents probably by removing the lower gate constriction (Staudner *et al*., 2024; Wang *et al*., 2019). In the present study we could reproduce this GOF effect. Indeed, in oocytes expressing the GOF PC2 construct (PC2 L677A N681A, further referred to as **PC2 AA**) large Na^+^ inward currents were observed under divalent cation free conditions (Fig. 1C, F). Replacing Na^+^ with NMDG^+^ in the bath solution completely blocked these inward currents. Similar to the inhibitory effect of PC1 on PC2 currents (Fig. 1EV), the PC2 AA mediated currents were significantly reduced by co-expressing PC1 (**PC2 AA+PC1**; Fig. 1D, F). This effect was not due to reduced cell surface expression of PC2 AA in the presence of PC1 (Fig. 1H; Appendix Fig.S2). Instead, it probably resulted from the formation of PC2 AA/PC1 heteromeric complexes as suggested by our co-immunoprecipitation experiments (Fig. 1I; Appendix Fig.S3, S4). In accordance with the structural data, three positively charged residues of PC1 (R4100, R4107, and H4111) may partially occlude the putative lower gate of the heteromeric channel thereby hindering the ion passage through the pore (Fig. 1D). Importantly, alanine substitutions of these PC1 residues (R4100A, R4107A, and H4111A) produced a novel GOF PC1 construct (further referred to as **PC1 AAA**), which formed complexes with PC2 AA (Fig. 1I) and dramatically increased whole-cell Na^+^ inward currents (**PC2 AA+PC1 AAA**; Fig. 1E, F). Neither PC1 nor PC1 AAA expressed alone elicited measurable currents, despite their trafficking to the plasma membrane (Appendix Fig.S5). This is consistent with the interpretation, that PC1 is unable to form functional homomeric ion channels, but can associate with PC2 to produce heteromeric ion channels. In additional experiments, we demonstrated that gradual substitution of PC1 by PC1 AAA in co-expression experiments with PC2 AA gradually increased the magnitude of Na^+^ inward currents (Appendix Fig.S6-A). This further confirmed the GOF effect of the alanine substitutions in PC1.

**Fig. 1.**
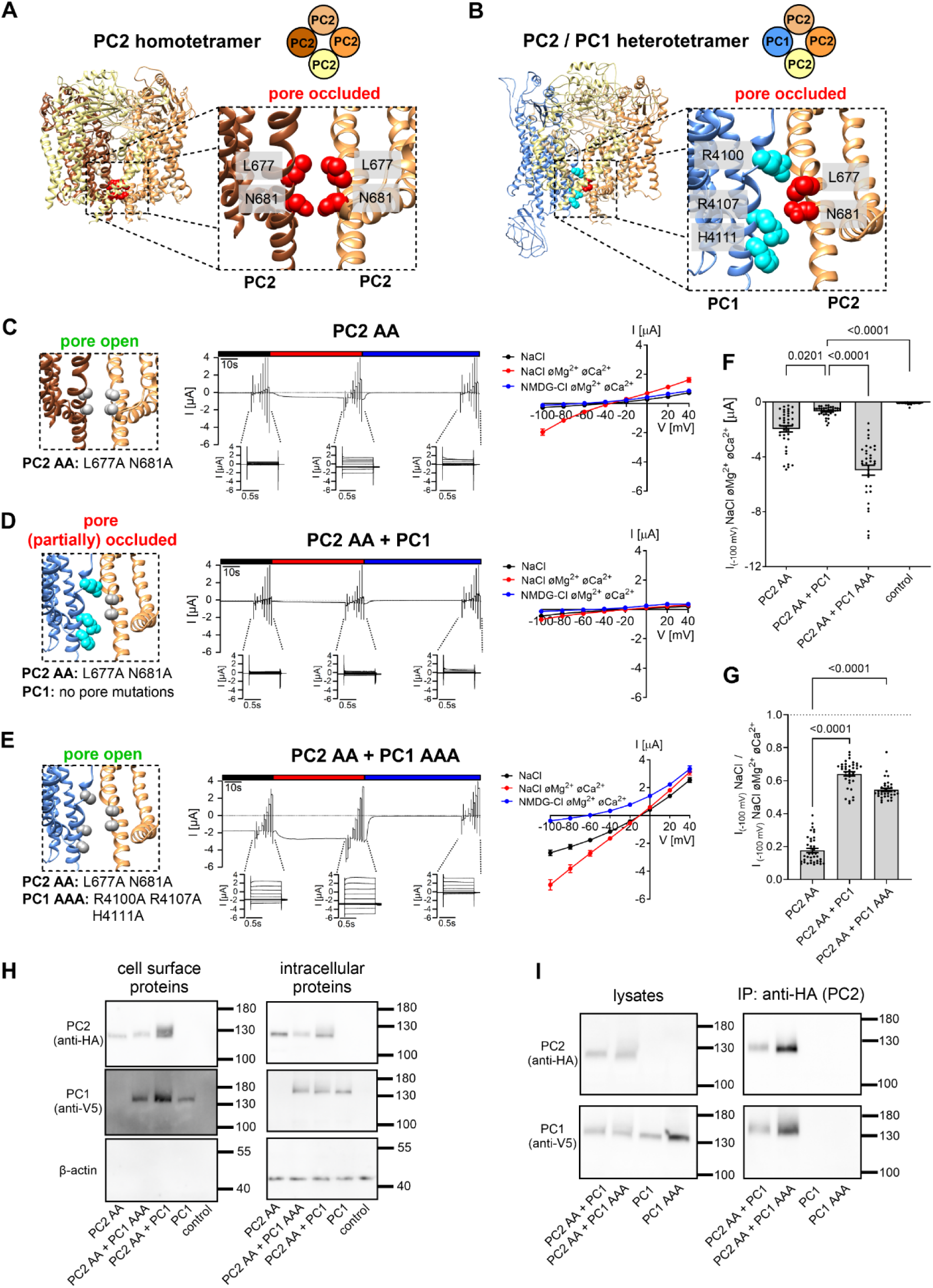
Generation of a gain-of-function (GOF) PC2/PC1 ion channel. ***A, B*** Side view of human homotetrameric PC2 (***A***) or heterotetrameric PC2/PC1 complex (***B***) in ribbon representation generated using atom coordinates from PDB entry 6T9N (Wang *et al*., 2020b) or 6A70 (Su *et al*., 2018a), respectively. Individual protomers are colored according to the schematic diagram shown above the structure. The insets show a portion of PC2 or PC2/PC1 on an expanded scale. Pore blocking residues are labeled and colored in *red* for PC2 and in *cyan* for PC1 with atoms shown as spheres. ***C-E*** *Left panels* Localization of alanine substitutions in grey within the pore of homomeric PC2 (***C***) or heteromeric PC2/PC1 ion channels (***D, E***) introduced to create PC2 AA and PC1 AAA GOF constructs. *Middle panels* Representative whole-cell current traces obtained in individual oocytes injected with 2.5 ng cRNA encoding human PC2 AA alone (***C***), or with additional co-injection of 5 ng cRNA encoding PC1 (***D***) or PC1 AAA (***E***). Presence of standard NaCl bath solution with or without divalent cations (øMg^2+^øCa^2+^) or NMDG-Cl bath solution without divalent cations is indicated by black, red, and blue bars, respectively. Overlays of whole-cell current traces resulting from voltage step protocols are shown below the continuous current recordings. *Right panels* Average I/V-plots (mean ± SEM) were constructed from similar recordings as shown in *middle panels* (***C***: *n* =39, N=4; ***D***: *n* =37, N=4; ***E***: *n* =34, N=4). ***F***, Maximal currents in PC2 AA *vs.* PC2 AA/PC1 *vs.* PC2 AA/PC1 AAA expressing oocytes. The maximal inward currents reached during the application of hyperpolarizing pulses of-100 mV in divalent free NaCl (NaCl øMg^2+^øCa^2+^) bath solution are shown. Data are from the same experiments as summarized in the I/V plots (***C***-***E***) and from control experiments (control) using oocytes solely injected with AS Cx38 (*n* =38, N=4). The *p*-values were calculated by the Kruskal-Wallis test with Dunn’s post hoc test. ***G***, Summary of the relative inhibitory effects of Ca^2+^ and Mg^2+^ on PC2 AA-, PC2 AA/PC1-and PC2 AA/PC1 AAA-mediated sodium inward currents (same experiments as in ***C***-***E***). In each individual recording, the maximal inward current reached in NaCl bath solution with divalent cations at-100 mV was normalized to the maximal inward current measured in NaCl bath solution without divalent cations at-100 mV. The *p*-values were calculated by the Kruskal-Wallis test with Dunn’s post hoc test. ***H***, Representative western blot analysis of cell surface (*left panels*) and intracellular (*right panels*) expression of PC2 and PC1 constructs in oocytes from one batch. Similar results were obtained in two other batches (N=3; Appendix Fig. S2). ***I***, Validation of PC2/PC1 complex formation using co-IP and PC2-HA as a “bait” protein. PC2 and PC1 were detected using western blot in co-IP preparations (*right panels*) and in corresponding cell lysates (*left panels*). PC2/PC1 complexes were isolated using an anti-HA antibody conjugated to magnetic beads, which recognized HA-tagged PC2. Original uncropped images of the same western blots are shown in Appendix Fig.S3.

We also noticed that the relative inhibitory effect of extracellular divalent cations on Na^+^ inward currents was significantly reduced in PC2 AA+PC1 and PC2 AA+PC1 AAA expressing oocytes compared to oocytes expressing PC2 AA alone (Fig. 1G). PC1 or PC1 AAA co-expression reduced divalent cation sensitivity to a similar extent. This indicates that the effect is independent of the GOF alanine mutations in PC1 (Appendix Fig.S6-B). Divalent cations are believed to block the ion permeation pathway of PC2 at least in part due to binding to the negatively charged D643 residues within the channel’s selectivity filter (Staudner *et al*., 2024; Wilkes *et al*., 2017). Thus, in PC2/PC1 heteromeric channels coordination of Ca^2+^ and Mg^2+^ by D643 is probably disturbed due to incorporation of PC1 into the pore.

In further experiments, we systematically investigated individual contributions of the R4100A, R4107A and H4111A substitutions in PC1 to the GOF effect (Fig. 2EV). Taken together, these results indicated that the R4107A substitution in PC1 was mainly responsible for the GOF effect, followed by the R4100A mutation. In contrast, the contribution of the H4111 residue to the pore occlusion appeared to be minor. Consistent with the results described above (Fig. 1G; Appendix Fig.S6-B), the inhibitory effect of divalent cations was reduced in all groups of oocytes co-expressing PC2 AA with different PC1 mutants (Fig. 2EV-C), compared to oocytes expressing PC2 AA alone (Fig. 1G). This indicated preferential formation of heteromeric PC2/PC1 ion channels in these oocytes. Furthermore, we demonstrated that co-expression of PC1 AAA with WT PC2 did not result in measurable inward currents (Fig. 2EV-A, B). It is therefore likely that both polycystins-PC1 and PC2-need to change their conformation from closed to open to produce fully active PC2/PC1 heteromeric channels. Finally, we demonstrated that the L677A mutation was more important for the GOF effect than the N681A mutation (Fig. 2EV-A, B).

In addition, we confirmed the dominant-negative effect of PC1 on another GOF PC2 construct (F604P) described previously (Wang *et al*., 2019). Indeed, the ion channel activity of PC2 F604P was completely blocked by either PC1 AAA or PC1 co-expression, suggesting formation of non-functional heteromeric PC2 F604P/PC1 ion channels (Appendix Fig.S7). The F604P mutation was shown to activate a specific gating mechanism of homomeric PC2, namely a π-α helical switch within the S6 transmembrane domain, which led to the opening of the channel’s lower gate (Zheng *et al*., 2018). This gating mechanism does not seem to function in PC2/PC1 heteromeric channels. Thus, it is tempting to speculate, that homomeric PC2 and heteromeric PC2/PC1 ion channels have distinct gating mechanisms. In contrast to our findings in the oocyte expression system, a stimulatory effect of PC1 on PC2 F604P has recently been described in HEK293 cells (Ha *et al*., 2024; Ha *et al*., 2020). We have no explanation for this, but it is conceivable that the function of PC2 and PC2/PC1 may vary in different expression systems.

In summary, we functionally validated a pore blocking effect of positively charged PC1 residues as predicted from structural analysis. In combination with the established PC2 AA GOF construct, alanine substitutions of these residues (R4100, R4107, H4111) in PC1 lead to a novel PC2 AA/PC1 AAA GOF construct.

### Heteromeric PC2/PC1 GOF ion channels exhibited significantly altered cation selectivity and reduced Ca^2+^ permeability compared to PC2 GOF channels

Using ion substitution experiments we compared the cation selectivity of heteromeric PC2 AA/PC1 AAA GOF ion channels with that of PC2 AA homomers (Fig. 2). Homomeric PC2 AA conducted K^+^ better than Na^+^, and Na^+^ slightly better than Li^+^ with no measurable conductance for NMDG^+^ (K^+^ > Na^+^ ≥ Li^+^ >>> NMDG^+^; Fig. 2A) consistent with our previous report (Staudner *et al*., 2024). In contrast, in oocytes co-expressing PC2 AA and PC1 AAA the measured inward currents were similar with Na^+^, Li^+^ or K^+^ as predominant cation in the bath solution. Thus, unlike PC2 AA, the heteromeric PC2 AA/PC1 AAA ion channel conducts K^+^, Na^+^ and Li^+^ equally well (K^+^ ≈ Na^+^ ≈ Li^+^ >>> NMDG^+^; Fig. 2B). In agreement with this, the reversal potentials in Na^+^ or Li^+^ containing bath solutions were close to zero in PC2 AA+PC1 AAA expressing oocytes, but considerably more negative in PC2 AA expressing oocytes. This also supports the conclusion that co-expression of PC2 AA with PC1 AAA results in ion channels which unlike PC2 AA homomeric channels have no preference for conducting K^+^ over Na^+^ or Li^+^. In addition, we calculated mean reversal potential shifts caused by replacing extracellular Na^+^ by K^+^, Li^+^ or NMDG^+^ to estimate cation permeability ratios (Table 1A). These estimates were in good agreement with the qualitative conclusions reached from the inward current data.

**Fig. 2.**
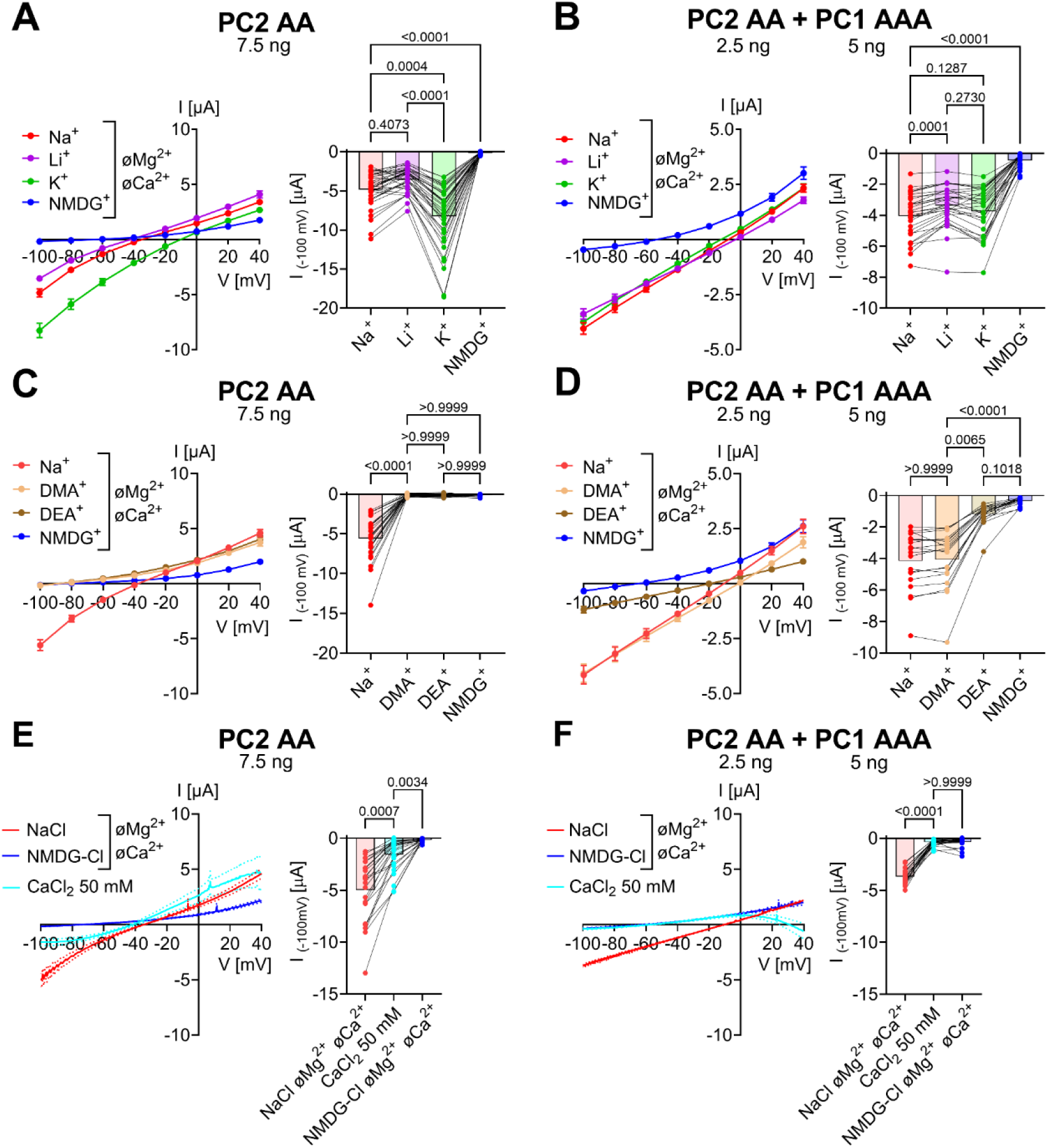
Homomeric PC2 AA and heteromeric PC2 AA / PC1 AAA ion channels demonstrate distinct cation permeability properties. ***A***-**D**, Permeability for small inorganic monovalent cations (***A*, *B***) and mid-size organic monovalent cations (***C*, *D***) was assessed for PC2 AA homomers (***A*, *C***) and PC2 AA / PC1 AAA heteromers (***B*, *D***) by replacing Na^+^ in the bath solution by Li^+^, K^+^, DMA^+^, DEA^+^ or NMDG^+^ in the absence of divalent cations. The channels’ permeability for Ca^2+^ (***E*, *F***) was estimated by applying 50 mM CaCl_2_ bath solution. *Left panels* Average I/V-plots (mean ± SEM) were obtained using a similar experimental approach as shown in Fig.1. To correct for endogenous oocyte currents, the average whole-cell current values measured in control oocytes (Appendix Fig. S8) were subtracted from the corresponding individual whole-cell current values measured in oocytes from the same batch expressing PC2 AA or co-expressing PC2 AA + PC1 AAA. *Right panels* Summary data show maximal inward currents reached during application of hyperpolarizing pulses of-100 mV in the presence of different cations in the bath as indicated. Average values and individual data points are shown (***A***, *n* =36, N=3; ***B***, *n* =30, N=3; ***C***, *n* =27, N=3; ***D***, *n* =19, N=3; ***E***, *n* =25, N=3; ***F***, *n* =25, N=3). Lines connect data points obtained from one oocyte. The *p*-values were calculated by the Friedman test with Dunn’s post hoc test.

**Table 1.** Effect of replacing Na^+^ in the bath solution by other small inorganic cations (A), mid-sized organic cations (B) or a large organic cation NMDG^+^ (A and B) on average reversal potentials in *Xenopus laevis* oocytes expressing PC2 AA alone or co-expressing PC2 AA with PC1 AAA or PC1. Reversal potentials (in mV) observed in the presence of Na^+^ (E_rev,Na_ ^+^), Li^+^ (E_rev,Li_^+^), K^+^ (E_rev,K_^+^), DMA^+^ (E_rev,DMA_^+^), DEA^+^ (E_rev,DEA_^+^) or NMDG^+^ (E_rev,NMDG_ ^+^) were estimated from the averaged *I*/*V*-curves shown in Fig. 2. E_rev,Na_ ^+^ was subtracted from E ^+^ measured in the bath solution containing cation X^+^ (Li^+^, K^+^, DMA^+^, DEA^+^ or NMDG^+^) rev,X instead of Na^+^ to calculate the corresponding reversal potential shift (ΔE_rev,X_ ^+^-Na ^+^). The reversal potential shifts were used to estimate relative permeability ratios (P_X_/P_Na_; see equation 1).

**(A) Relative permeability ratios for $\text{Na}^+$ , $\text{Li}^+$ , $\text{K}^+$**
|  | **PC2 AA** | **PC2 AA+PC1 AAA** | **PC2 AA+PC1** |
| --- | --- | --- | --- |
| $E_{\text{rev},\text{Na}^+}$ [mV] | - 35 | - 8 | - 13 |
| $E_{\text{rev},\text{Li}^+}$ [mV] | - 42 | - 3 | - 8 |
| $E_{\text{rev},\text{K}^+}$ [mV] | - 10 | - 13 | - 15 |
| $E_{\text{rev},\text{NMDG}^+}$ [mV] | - 65 | - 52 | - 58 |
| $\Delta E_{\text{rev},\text{Li}^+-\text{Na}^+}$ [mV] | - 7 | 5 | 5 |
| $\Delta E_{\text{rev},\text{K}^+-\text{Na}^+}$ [mV] | 25 | - 5 | - 2 |
| $\Delta E_{\text{rev},\text{NMDG}^+-\text{Na}^+}$ [mV] | - 30 | - 44 | - 45 |
| $P_{\text{Na}} : P_{\text{Li}} : P_{\text{K}} : P_{\text{NMDG}}$ | 1.0 : 0.8 : 2.7 : 0.3 | 1.0 : 1.2 : 0.8 : 0.2 | 1.0 : 1.2 : 0.9 : 0.2 |

**(B) Relative permeability ratios for $\text{DMA}^+$ , $\text{DEA}^+$ and NMDG $^+$**
|  | **PC2 AA** | **PC2 AA+PC1 AAA** | **PC2 AA+PC1** |
| --- | --- | --- | --- |
| $E_{\text{rev},\text{Na}^+}$ [mV] | - 38 | - 10 | - 15 |
| $E_{\text{rev},\text{DMA}^+}$ [mV] | - 87 | - 1 | - 18 |
| $E_{\text{rev},\text{DEA}^+}$ [mV] | - 89 | - 20 | - 46 |
| $E_{\text{rev},\text{NMDG}^+}$ [mV] | - 84 | - 65 | - 71 |
| $\Delta E_{\text{rev},\text{DMA}^+-\text{Na}^+}$ [mV] | - 49 | 9 | - 3 |
| $\Delta E_{\text{rev},\text{DEA}^+-\text{Na}^+}$ [mV] | - 51 | - 10 | - 31 |
| $\Delta E_{\text{rev},\text{NMDG}^+-\text{Na}^+}$ [mV] | - 46 | - 55 | - 56 |
| $P_{\text{Na}} : P_{\text{DMA}} : P_{\text{DEA}} : P_{\text{NMDG}}$ | 1.0 : 0.1 : 0.1 : 0.2 | 1.0 : 1.4 : 0.7 : 0.1 | 1.0 : 0.9 : 0.3 : 0.1 |

Furthermore, we investigated permeability of homomeric PC2 AA *versus* heteromeric PC2 AA/PC1 AAA ion channels for mid-sized organic monovalent cations (DMA^+^, DEA^+^). Homomeric PC2 GOF channels were almost impermeable for these organic cations (Na^+^ >>> DMA^+^ ≈ DEA^+^ ≈ NMDG^+^; Fig. 2C; Table 1B). In contrast, heteromeric PC2/PC1 GOF channels conducted DMA^+^ and Na^+^ equally well and DEA^+^ significantly better than NMDG^+^ (Na^+^ ≈ DMA^+^ > DEA^+^ > NMDG^+^; Fig. 2D; Table 1B). Similar cation permeability properties were observed in oocytes co-expressing PC2 AA and PC1 without alanine substitutions (PC2 AA+PC1; Fig. 3EV; Table 1A and B), consistent with previous findings (Wang *et al*., 2019). In our experiments the amount of injected cRNA was increased by 3-fold for both polycystins (PC2AA and PC1) to achieve higher currents, thereby improving the resolution of the current measurements. From this electrophysiological analysis we concluded, that heteromerization of PC2 AA with PC1 AAA or PC1 resulted in formation of ion channels with significantly altered ion channel properties compared to PC2 homomers. Importantly, this effect was not due to GOF alanine substitutions in the putative lower gate of PC1. Instead, it can be attributed to structural changes of the channel’s selectivity filter induced by PC1.

Finally, we assessed Ca^2+^ permeability of PC2 AA and PC2 AA / PC1 AAA by applying 50 mM CaCl_2_ bath solution according to a previously established protocol exploiting endogenous Ca^2+^-activated Cl^-^ channels for the detection of Ca^2+^ influx (Grosch *et al*., 2021; Staudner *et al*., 2024). Consistent with our previous observation (Staudner *et al*., 2024), in the presence of 50 mM CaCl_2_ we observed substantial inward currents in PC2 AA expressing oocytes (Fig. 2E). In contrast, this was not the case in oocytes co-expressing PC2 AA and PC1 AAA (Fig. 2F). This finding indicates that PC2 AA is permeable for Ca^2+^, whereas the Ca^2+^ permeability of PC2 AA/PC1 AAA heteromers is abolished or below the detection limit of our assay. We also did not detect any permeability for Mg^2+^ or Ba^2+^ in oocytes co-expressing PC2 AA and PC1 AAA (Appendix Fig. S9).

Next, we slightly modified our experimental protocol to enhance Ca^2+^ entry into oocytes by reducing the voltage-dependent pore blocking effect of extracellular Ca^2+^ (Arif Pavel *et al*., 2016; Wang *et al*., 2019). Indeed, this modification resulted in a significantly stronger Ca^2+^-mediated current response in PC2 AA expressing oocytes (Appendix Fig. S10A). However, under the same conditions only a marginal Ca^2+^ permeability could be revealed in PC2 AA + PC1 AAA expressing oocytes (Appendix Fig. S10B). Interestingly, another study comparing the Ca^2+^ permeability of PC2 AA with that of PC2 AA / PC1 heteromeric ion channels came to the opposite conclusion (Wang *et al*., 2019). At present, we do not have an explanation for these conflicting results, but it is noteworthy that in the latter study the amount of cRNA injected was very high (30 ng/oocyte for PC2 and 60 ng/oocyte for PC1 compared to 2.5 ng/oocyte for PC2 and 5 ng/oocyte for PC1 in our study). Taken together, our findings suggest that PC2 homomers and PC2/PC1 heteromers are two distinct types of ion channels probably with different (patho-)physiological roles in different tissues and subcellular localizations. This seems plausible, because PC2 and PC1 demonstrate profound differences regarding their spatiotemporal expression pattern in kidney and other organs (Cantero & Cantiello, 2022; Chauvet *et al*, 2002; Geng *et al*, 1996; Markowitz *et al*, 1999; Ong *et al*, 1999).

### PC2 preferentially forms heteromeric complexes with PC1

When PC1 and PC2 are co-expressed, formation of heteromeric PC2/PC1 and homomeric PC2 complexes may occur (Fig. 3A). To determine whether heteromers or homomers are preferentially assembled in the oocyte expression system, we performed “PC1 titration” experiments by co-expressing a fixed amount of PC2 AA with increasing amounts of PC1 AAA. Using the same experimental protocol as shown in Fig. 1, we estimated the average reversal potential in NaCl divalent cation free bath solution in each group of oocytes (Fig. 3B, C; Appendix Fig. S11). Increasing the PC1 amount resulted in a progressive shift of the reversal potential from approximately-35 mV to-10 mV. This can be explained by an increased proportion of heteromeric PC2 AA/PC1 AAA channels at the cell surface, because the heteromeric channels have a reduced potassium to sodium permeability ratio compared to homomeric PC2 AA channels, in accordance with the results shown in Fig. 2 A, B. Importantly, considerable heteromerization was evident even with the smallest amount of injected PC1 AAA cRNA (0.1 ng). Moreover, heteromerization reached saturation when 1 ng of PC1 AAA was co-expressed with 2.5 ng of PC2 AA. This corresponds to 0.77 pmol and 2.5 pmol of injected cRNA encoding PC1 AAA and PC2 AA, respectively. Thus, assuming a similar translation efficiency of the cRNAs, we can estimate a PC2/PC1 molar ratio of 3.2:1, at which the apparent saturation of heteromerization was reached. This estimate is in good agreement with the 3:1 PC1/PC2 stoichiometry observed in the cryo-EM structure (Su *et al*., 2018a) and with previous biochemical and biophysical results (Yu *et al*., 2009). Furthermore, data from the same experiments were re-analyzed to calculate the relative inhibitory effect of Ca^2+^ on sodium inward currents in each group of oocytes (Fig. 3D). As shown in Fig. 1G, homomeric PC2 AA channels demonstrated significantly higher sensitivity to extracellular Ca^2+^ compared to heteromeric PC2 AA/PC1 AAA channels. Therefore, in addition to the reversal potential shift, the reduction of Ca^2+^ sensitivity can be used as a parameter to estimate the dependence of the PC2/PC1 heteromerization process on PC1 dosage. Indeed, both parameters demonstrated essentially the same dependence on PC1 AAA dosage (Fig. 3D). Taken together, these results demonstrate that PC2 preferentially forms heteromeric complexes when co-expressed with PC1 most likely with a 3:1 stoichiometry.

**Fig. 3.**
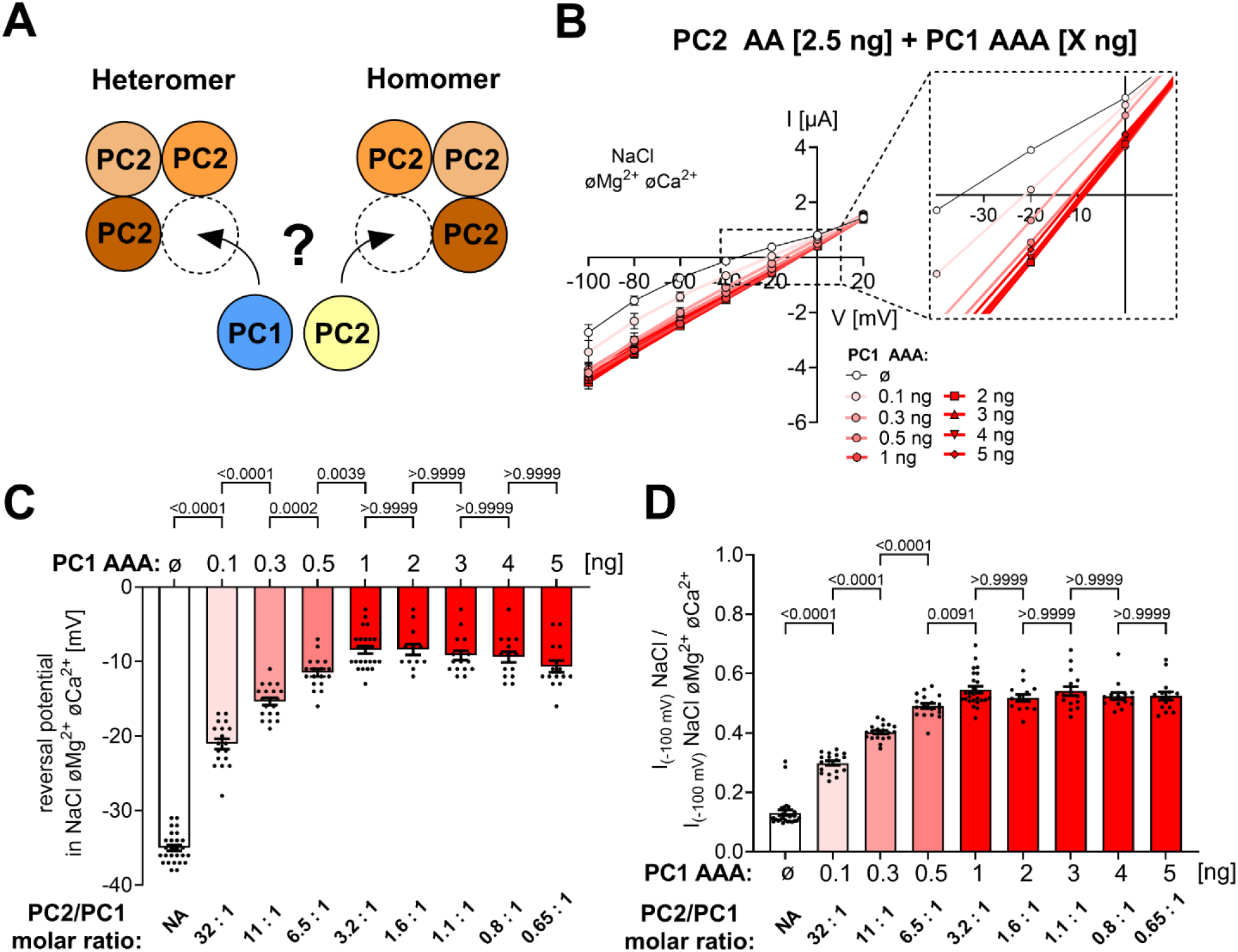
PC2 preferentially forms heterotetrameric complexes with PC1 probably in a 3:1 stoichiometry. ***A*,** Schematic diagram illustrating two possibilities of PC1 and PC2 oligomerization. ***B***, Average I/V-plots (mean ± SEM) obtained in NaCl bath solution without divalent cations (NaCl øMg^2+^øCa^2+^) from oocytes co-injected with a constant amount of PC2 AA (2.5 ng) and a variable amount of PC1 AAA as indicated. The inset showing a portion of the same I/V-plots on an expanded scale demonstrates a progressive shift of the reversal potential with increasing amount of PC1 AAA. This effect saturates at about 1 ng of injected PC1 AAA. Complete set of I/V-plots obtained from these oocytes are shown in Appendix Fig.S11. ***C***, Summary of the reversal potentials from the same experiments as shown in (***B***). ***D***, Summary of the relative inhibitory effects of Ca^2+^ and Mg^2+^ on PC2AA-and PC2 AA/PC1 AAA-mediated sodium inward currents calculated as described in Fig. 1G. Mean ± SEM and values from individual recordings are shown (13≤*n*≤27, N=2-3). The *p*-values were calculated by the one-way ANOVA with Bonferroni’s post hoc test.

### Pore-forming domain of PC1 probably exhibits a canonical TRP-like conformation

The published structure of the PC2/PC1 heterotetrameric complex suggests that the PC1 subunit has a noncanonical architecture of the pore-forming domains (Su *et al*., 2018a). Unlike TRP channels, such as PC2 homomers (Grieben *et al*., 2017; Wilkes *et al*., 2017), PC2L1 homomers (Hulse *et al*, 2018; Su *et al*, 2018b), and PC2L1/PC1L3 heteromers (Su *et al*, 2021), it lacks the two pore loop helices (PH1 and PH2). Interestingly, the proximal portion of the pore loop (C4051-W4080) was not resolved in the original model (Fig. 4A, *left structure*; PDB ID: 6A70). Additionally, the distal part of the pore loop, which resembled pore helix 1 (PH1), directly transitions into the S6 transmembrane domain and was thus interpreted as the extracellular portion of the S6 helix-S6a (Fig. 4A). We have re-interpreted the original cryo-EM volume (EMD-6991) and generated an alternative model of the PC2/PC1 complex that differs significantly from the initial interpretation in certain aspects. Model-versus-data comparisons indicate that the new model aligns with the original cryo-EM volume more accurately than the previous model does (Appendix Table S1).

**Fig. 4.**
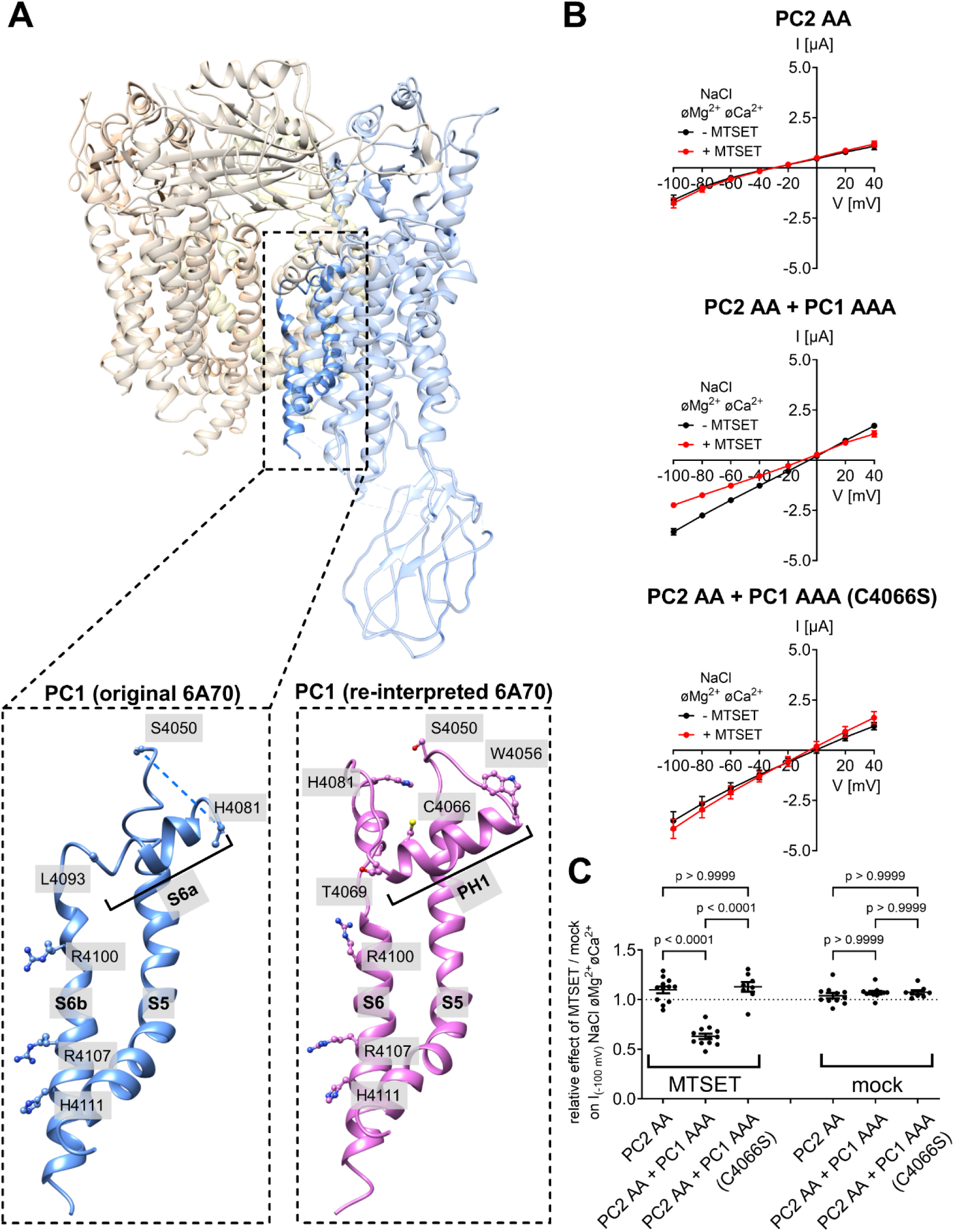
The pore domain of PC1 probably exhibits a canonical TRP-like conformation. ***A*,** Side view of human heterotetrameric PC2/PC1 complex in ribbon representation generated using the published atom coordinates from PDB entry 6A70 (Su *et al*., 2018a). Individual protomers are colored similarly as in Fig. 1B. The inset shows the pore domain of PC1 on an expanded scale. The corresponding cryo-EM data was re-interpreted (see Fig. 4EV) to generate a new model of the PC1 pore domain shown to the right of the published model. PH1, pore helix 1. ***B***, Validation of the new PC1 model using MTSET treatment and electrophysiological measurements. Average I/V-plots (mean ± SEM) obtained in the NaCl bath solution without divalent cations (NaCl øMg^2+^øCa^2+^) from oocytes expressing PC2 AA alone, co-expressing PC2 AA and PC1 AAA, or co-expressing PC2 AA and PC1 AAA with additional C4066S mutation before (-MTSET) or after 3 min incubation in NaCl bath solution supplemented with 1 mM MTSET (+ MTSET). Complete set of I/V-plots obtained from these oocytes are shown in Appendix Fig.S12. The oocyte was unclamped during the incubation time. Before the second current measurement, MTSET was washed out with NaCl bath solution. Impaling microelectrodes were not removed from the oocyte until the end of the experiment. ***C***, Summary data (mean ± SEM and individual data points) from experiments shown in ***B*** and from similar experiments performed with mock-treated control oocytes shown in Appendix Fig.S12 are presented. Relative effects of MTSET / mock incubation on the maximal inward currents reached at-100 mV in divalent free NaCl (NaCl øMg^2+^øCa^2+^) bath solution were calculated. In each individual recording, the current measured after MTSET or mock incubation was normalized to the current measured before the treatment (8≤*n*≤12, N=2-3). The *p*-values were calculated by the one-way ANOVA with Bonferroni’s post hoc test.

We identified an unassigned cylindrical density in the pore forming region of the PC2/PC1 complex (Fig. 4EV-A-D, indicated by *red arrow*). This discovery led us to re-assess the EM densities of the pore-forming domains of PC1 by using a homology model of PC1, which included the S5-pore loop-S6 segments and was based on the PC2 structure (26.11% sequence identity). Refining this model against the published EM map suggested a plausible configuration of PC1, wherein a canonical TRP-like arrangement of the S5-PH1-S6 segments is feasible (Fig. 4A, *right structure*; Fig. 4EV-B, D, F). We interpreted the previously unassigned cylindrical density as part of the continuous S6 helix (H4081-V4088; Fig. 4EV-B, D, F), indicating that the S6 helix likely remains intact and is not divided into S6a and S6b sections, as initially proposed. Additionally, the more proximal sequence (W4056-T4069) with its medium-sized side chains aligns better with the putative PH1 EM density than the originally suggested H4081-L4093 sequence (Fig. 4EV-A, C, E).

We identified a density to accommodate the C4066 near the distal end of PH1 (Fig. 4EV-D; Fig. 4A, *right structure*). To confirm our structural prediction, we performed functional experiments using a sulfhydryl reagent MTSET to covalently modify the C4066 residue. In oocytes co-expressing PC2 AA and PC1 AAA, MTSET incubation led to a significant and irreversible decrease in currents (Fig. 4B, C; Appendix Fig.S12). This effect was not observed in mock control experiments or in oocytes expressing PC2 AA alone. Notably, the inhibitory effect of MTSET was absent in oocytes co-expressing PC2 AA and a C4066S mutant of PC1 AAA (Fig. 4B, C; Appendix Fig.S12). Application of DTT completely reversed the inhibitory effect of MTSET on PC2 AA/PC1 AAA channels (Fig. 5EV), likely by removing the adduct from the cysteine residue (Kellenberger *et al*, 2002). These results indicate that the pore loop of PC1 has a canonical TRP-like conformation.

In summary, our findings provide strong evidence that PC1 and PC2 can form heteromeric ion channels with ion channel properties different from those of PC2 homomeric channels. We validated the published structure of the PC2/PC1 complex by generating a novel GOF PC2/PC1 heteromeric ion channel (PC2 AA / PC1 AAA). This PC2/PC1 GOF channel may serve as a valuable tool to investigate the impact of ADPKD-associated mutations on the formation and cation conductance of heteromeric PC2/PC1 ion channel complexes. These future investigations may enhance our understanding of the role of PC2/PC1 ion channels in the pathogenesis of ADPKD and support the concept that ADPKD is a channelopathy (Grosch *et al*., 2021; Staudner *et al*., 2024; Vien *et al*, 2020; Wang *et al*., 2023).

## Methods

### Reagents and tools

See Table 2.

**Table 2.**
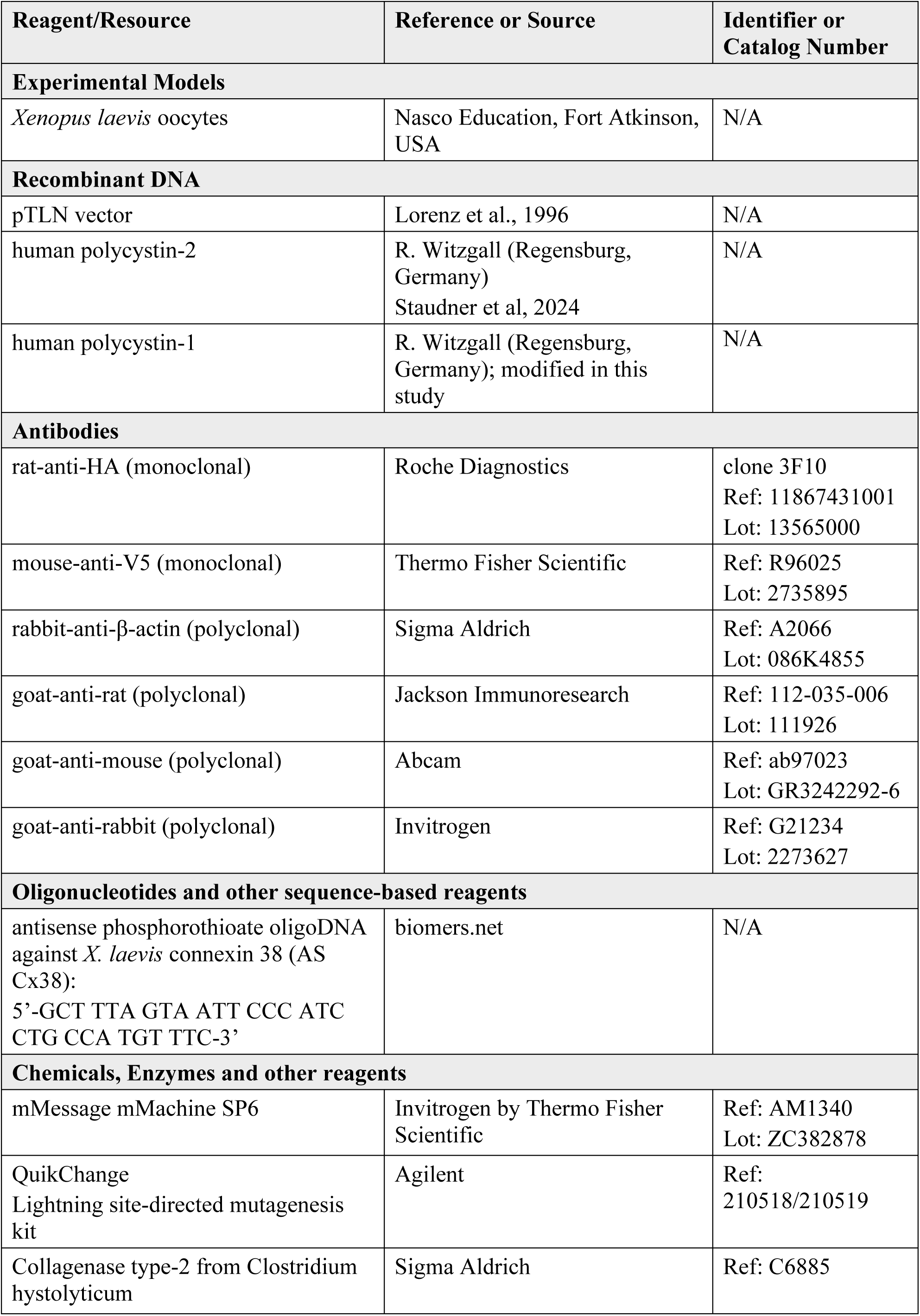
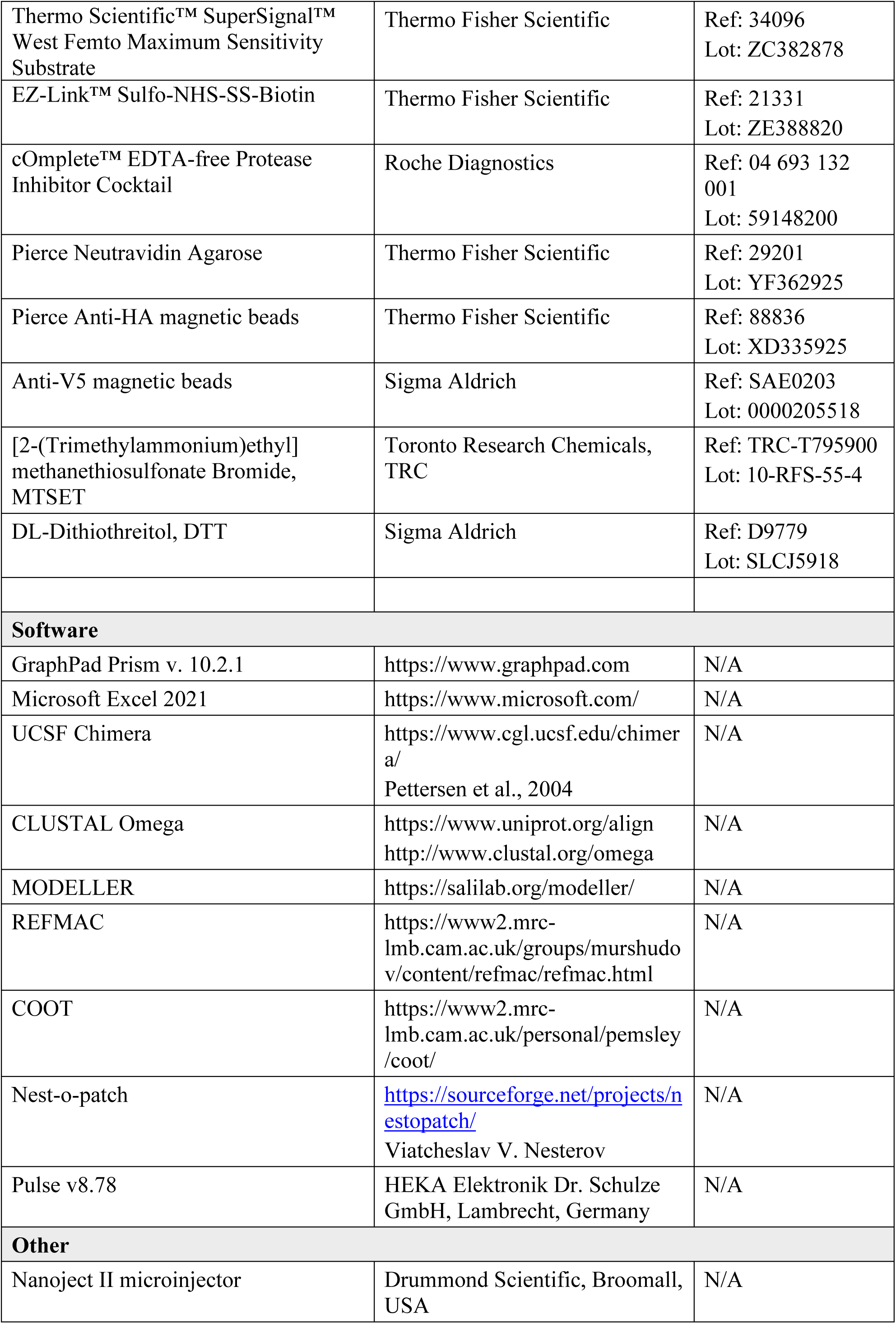
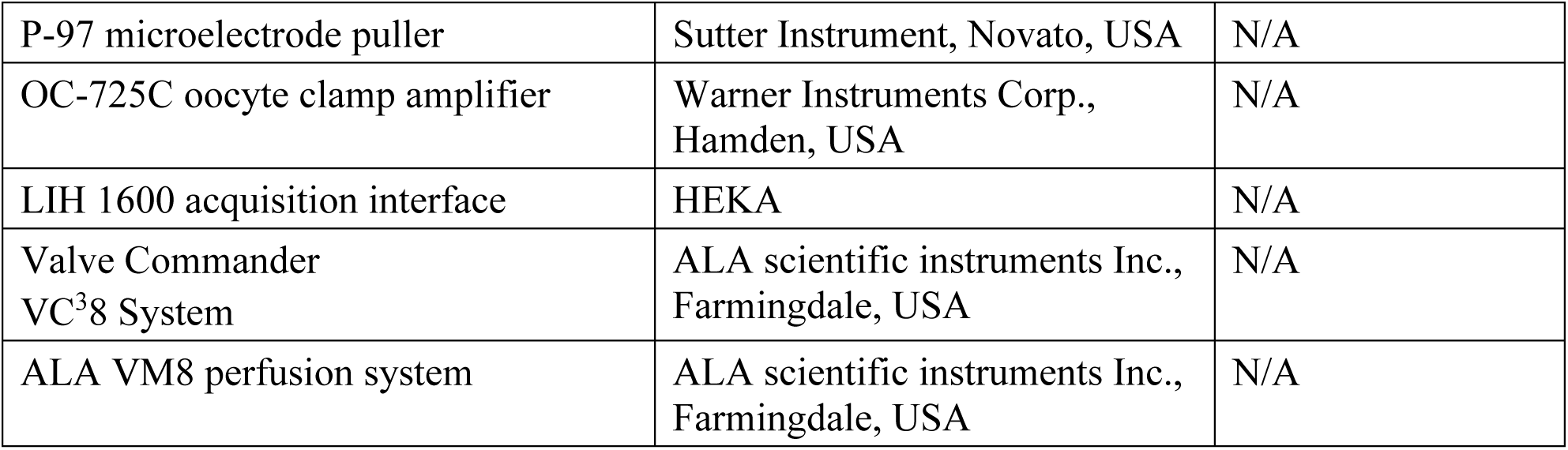
Reagents and Tools.

#### cDNA constructs

Full-length complementary DNAs (cDNAs) encoding human polycystin-1 and polycystin-2 were kindly provided by R. Witzgall (Regensburg, Germany). The first 3,048 amino acids of PC1 were removed to obtain the C-terminal fragment of PC1 (PC1-CTF, referred to as PC1 in the manuscript) using Gibson assembly. For heterologous expression in *Xenopus laevis* oocytes, PC1 and PC2 were subcloned into the pTLN vector (Lorenz *et al*, 1996). Mutations were introduced to PC1 and PC2 using the QuikChange Lightning site-directed mutagenesis kit (Agilent). For immunoblotting, PC1 and PC2 constructs were modified by attaching a C-terminal V5-tag (GKPIPNPLLGLDST) and an N-terminal HA-tag (YPYDVPDYA), respectively. In addition, an Ig k-chain leader sequence (METDTLLLWVLLLWVPGSTGD) was introduced to the N-terminus of PC1 to improve its cell surface expression (Wang *et al*., 2019). Sequences of all constructs were routinely verified by a commercially available sequence analysis service (LGC Genomics GmbH, Germany). Plasmids were linearized using the MluI restriction enzyme and used as templates for cRNA synthesis using SP6 RNA polymerase (mMessage mMachine SP6, Ambion).

#### Isolation of Xenopus laevis oocytes and two-electrode voltage-clamp (TEVC) experiments

Isolation of *Xenopus laevis* oocytes and two-electrode voltage-clamp (TEVC) experiments were performed essentially as described previously (Staudner *et al*., 2024; Sure *et al*, 2022). In brief, ovarian lobes were excised by partial ovariectomy under anaesthesia with Tricain 0.2%, in accordance with the principles of German legislation, with approval by the animal welfare officer for the University of Erlangen-Nürnberg (FAU), and under the governance of the state veterinary health inspectorate. The isolated sections of the ovarian lobes were then treated with 600-700 U/ml collagenase type-2 from Clostridium hystolyticum (Sigma-Aldrich) for 3-4 hours at 19°C. Subsequently, stage V-VI oocytes were injected with the amount of cRNA coding for the respective PC2 and/or PC1 construct as indicated in the figures or the figure legends. To avoid background currents mediated by endogenous connexin 38 hemichannels under divalent cation-free conditions, 3 ng of an antisense phosphorothioate DNA oligomer corresponding to nucleotides-5 to +25 relative to the coding region of connexin 38 (5’-GCTTTAGTAATTCCCATCCTGCCATGTTTC-3’; AS Cx38) was co-injected in every oocyte. Oocytes injected only with the AS Cx38 were used as control group. Following the cRNA injection, oocytes were incubated at 19°C for 48-72 hours in ND9: NaCl 9 mM, KCl 2 mM, N-methyl-D-glutamine-Cl (NMDG-Cl) 87 mM, CaCl_2_ 1.8 mM, MgCl_2_ 1 mM, HEPES 5 mM, pH 7.4 with Tris supplemented with 100 units/ml of sodium penicillin and 100 µg/ml of streptomycin sulphate.

In TEVC experiments conducted at room temperature, oocytes were continuously superfused with a bath solution as indicated in the figures. The standard NaCl bath solution was composed as follows: NaCl 96 mM, KCl 4mM, CaCl_2_ 1mM, MgCl_2_ 1mM, HEPES 10 mM, pH = 7.4 adjusted with Tris. CaCl_2_ and MgCl_2_ were excluded from the standard NaCl bath solution to create a NaCl bath solution nominally free of divalent cations (NaCl øMg^2+^ øCa^2+^). In the latter solution 95 mM NaCl were replaced with the same concentration of NMDG-Cl to obtain NMDG-Cl divalent cation-free bath solution (NMDG-Cl øMg^2+^ øCa^2+^). Gravity-based bath solution exchanges were controlled by a magnetic valve system (ALA VM8; ALA Scientific Instruments). The holding potential was constantly kept at-60 mV. At the end of each solution application, a voltage step protocol with consecutive 1,000 ms voltage steps in 20 mV increments starting with a hyperpolarizing pulse to-100 mV from a holding potential of-60 mV and ending with a depolarizing pulse to +40mV was performed. The current values measured during the final 300 ms of each pulse were used to construct the corresponding I/V-plot.

To estimate the channels’ selectivity for small inorganic monovalent cations (Na^+^, Li^+^, K^+^) and mid-sized organic monovalent cations (DMA^+^, DEA^+^) (Fig.2A-D; Fig. 3EV; Appendix Fig. S8A, B), bath solutions with the following compositions were used: NaCl, LiCl, KCl, DMA-Cl, DEA-Cl or NMDG-Cl 100 mM, HEPES 10 mM, pH 7.4 adjusted with Tris. To correct for endogenous oocyte conductances, average whole-cell current values obtained in the corresponding bath solution in control oocytes (Appendix Fig.S8) were subtracted from current values obtained in polycystin expressing oocytes from the same batch. The permeability ratios for monovalent cations (P_Na_: P_K_: P_Li_: P_DEA_: P_DMA_: P_NMDG_) were calculated from the corresponding reversal potential shifts using the modified GHK equation (Hille, 2001):

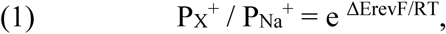

where ΔE_rev_ represents the shift of the reversal potential (E_rev_) caused by replacing Na^+^ in the bath solution with cation X^+^ [ΔE_rev_ = E_rev,X_^+^-E_rev,Na_ ^+^], *F* is the Faraday’s constant, *R* is the universal gas constant, and *T* is the absolute temperature in Kelvin.

To estimate the channels’ Ca^2+^ permeability, a bath solution with a high Ca^2+^ concentration (CaCl_2_ 50 mM, HEPES 10 mM, pH 7.4 adjusted with Tris) was applied for 60 s to oocytes continuously clamped at the standard holding potential of-60 mV (Fig.2E, F). Current-voltage (I/V) relationships were obtained using a 2-s voltage ramp protocol from-100 mV to +40 mV performed at the end of the CaCl_2_ application period. As previously reported, inward currents elicited under these conditions in oocytes expressing either PC2L1 or GOF PC2 (L677A/N681A) were suppressed by pre-injecting oocytes with 50 mM K-EGTA and could be attributed to Ca^2+^-activated Cl^-^ channels abundantly expressed in *Xenopus laevis* oocytes (Grosch *et al*., 2021; Staudner *et al*., 2024). Thus, with this experimental approach stimulation of Ca^2+^-activated Cl^-^ channels can be used as a surrogate parameter to assess polycystin-mediated Ca^2+^ influx. In some experiments with 50 mM CaCl_2_ in the bath solution a holding potential of 0 mV instead of-60 mV was used to reduce the known voltage-dependent blocking effect of Ca^2+^ on its own passage through the channel’s pore (Arif Pavel *et al*., 2016). Bath solutions containing 50 mM BaCl_2_ or 50 mM MgCl_2_ instead of 50 mM CaCl_2_ were used to assess the channels’ permeability for Ba^2+^ or Mg^2+^, respectively. Data analysis was performed using the programs Microsoft Excel and ‘Nest-o-Patch’ (http://sourceforge.net/projects/nestopatch) written by Dr. V. Nesterov (Friedrich-Alexander-Universität Erlangen-Nürnberg, Institute of Cellular and Molecular Physiology, Erlangen, Germany).

#### Detection of polycystin-1 and polycystin-2 at the cell surface

Cell surface expression of PC1 and PC2 was assessed using a biotinylation approach essentially as described previously (Ilyaskin *et al*, 2021; Staudner *et al*, 2024; Sure *et al*, 2022). In brief, oocytes were incubated with EZ-Link Sulfo-NHS-SS-Biotin (Thermo Fisher Scientific) and lysed mechanically through a 27G-needle. Subsequently, biotinylated proteins were separated from intracellular proteins with Pierce NeutrAvidin agarose beads (Thermo Fisher Scientific). After protein separation by electrophoresis, V5-tagged PC1 or HA-tagged PC2 was detected on a PVDF membrane. The primary antibodies used were a monoclonal rat anti-HA antibody (Roche Diagnostics, dilution: 1:1,000) and monoclonal mouse anti-V5 antibody (Thermo Fisher Scientific, dilution: 1:2,000). The secondary antibodies used were horseradish peroxidase-conjugated goat-anti-rat antibody (Jackson Immunoresearch, dilution: 1:10,000) and horseradish peroxidase-conjugated goat-anti-mouse (Abcam, dilution: 1:50,000). To confirm the separation of cell surface proteins from intracellular proteins, western blots were subsequently stripped and reanalysed using a polyclonal rabbit anti-β-actin antibody (Sigma-Aldrich, dilution: 1:5,000) and a secondary horseradish peroxidase-conjugated goat-anti-rabbit antibody (Invitrogen, dilution: 1:40,000).

#### Co-immunoprecipitation (co-IP)

30 oocytes per group were washed three times in ND9 on ice. Oocytes were subsequently transferred into 1 mL lysis buffer (Tris 25mM, NaCl 150 mM, EDTA 1mM, NP40 1%, Glycerol 5%, supplemented with Roche cOmplete™ protease inhibitor cocktail, pH = 7.4) and mechanically lysed using a syringe and 27G-needle. Lysates were incubated by rotating at 4°C for 1 hour. Lysates were then centrifuged for 30 minutes at 10,000 g at 4°C and supernatants were collected. Before application to the lysates, anti-HA (Thermo Fisher Scientific) or anti-V5 (Sigma Aldrich) antibody-coated magnetic beads were washed two times in TBST (Tris 25mM, NaCl 150mM, TWEEN 0.05%, pH = 7.5). 25 µL of the respective beads were pipetted on the collected supernatant and subsequently incubated for 30 minutes at room temperature by rotation. Thereafter, beads were collected and washed three times in TBST and once in nuclease-free water for 5 minutes at 4°C. For elution of the precipitated proteins the beads were incubated for 10 minutes in 100 µL of elution buffer (0.1M glycine, pH = 2) at room temperature. After removal of the magnetic beads, eluted protein fractions were neutralized with 15 µL of neutralization buffer (1M Tris, pH = 8.5).

#### Re-analysis of cryo-electron microscopy data and modelling approaches

The structure of the human PC2/PC1 heterotetrameric complex was retrieved from the RCSB protein database (PDB ID: 6A70;(Su *et al*., 2018a)). and the corresponding EM density was obtained from EMDB (*Electron Microscopy Data Bank*; EMD-6991). To generate a homology model of the PC1 C-terminal domain (CTD), sequence alignment between the template structure (PDB ID: 5MKF;(Wilkes *et al*., 2017)) and the target sequence of PC1 CTD was performed using the CLUSTAL Omega multiple sequence alignment tool available at www.uniprot.org/align. MODELLER (Sali & Blundell, 1993) was then employed for comparative structure modeling to generate an initial model of the PC1 CTD, which was subsequently fitted into the corresponding EM density (EMD-6991). Following manual corrections to the sequence alignment, the final model underwent iterative refinement using REFMAC (Murshudov *et al*, 1997) against the EM density. Regions that could not be modeled based on homology were either manually adjusted or built *de novo* using COOT (Emsley *et al*, 2010).

#### Statistical Analysis and Data presentation

N indicates the number of different batches of *Xenopus laevis* oocytes, and *n* indicates the number of individual oocytes analysed per experimental group. Data are presented as mean ± SEM. Statistical evaluation was done using an appropriate statistical test as indicated in figure legends using GraphPad Prism (GraphPad Software Inc.). The normal distribution of data was assessed using the D’Agostino-Pearson omnibus test. Graphical representations were created using GraphPad Prism. Structure visualizations were prepared using UCSF Chimera developed by the Resource for Biocomputing, Visualization, and Informatics at the University of California, San Francisco, with support from the National Institutes of Health P41-GM103311 (Pettersen *et al*, 2004).

## Data Availability

This study includes no data deposited in external repositories.

## Acknowledgements

We gratefully acknowledge the expert technical assistance of Ralf Rinke, Sonja Mayer, and Jessica Rinke. Funded by the Interdisziplinäres Zentrum für Klinische Forschung (IZKF, Interdisciplinary Center for Clinical Research Erlangen; subproject F8 to C.K.; MD-Thesis scholarships to T.S. and L.G.) and the Deutsche Forschungsgemeinschaft (DFG, German Research Foundation) – Projektnummer 517598260 (to A.I.).

## Disclosure and competing interests statement

The authors declare that they have no conflicts of interest with the contents of this article.

## Expanded View Figures and Figure legends

**Fig. 1EV.**
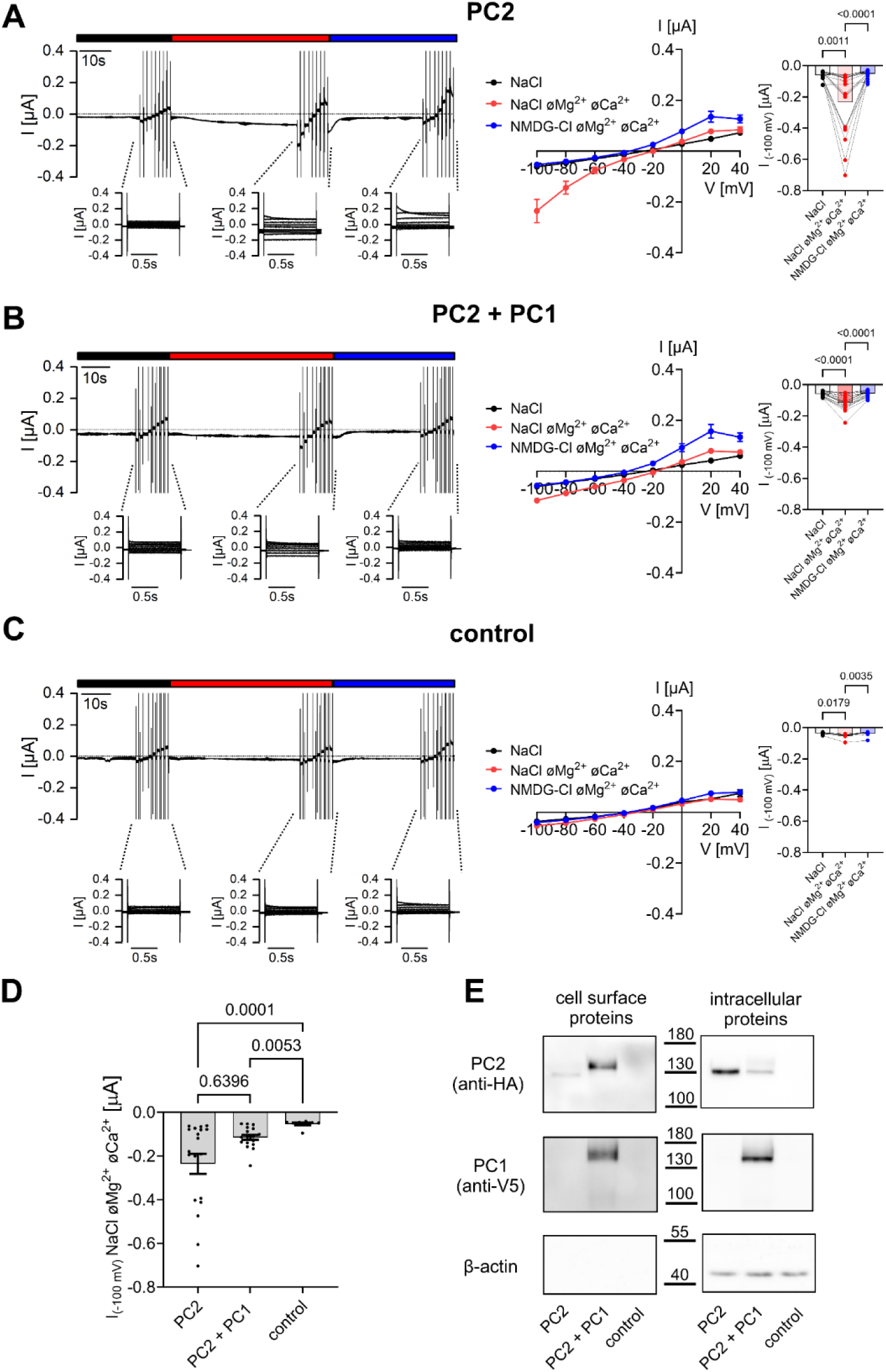
Ion channel activity of PC2 expressed alone or together with PC1 is low. ***A-C*** *Left panels* Representative whole-cell current trace obtained in an oocyte injected with 7.5 ng cRNA encoding human PC2 alone (***A***), or with additional co-injection of 15 ng cRNA encoding PC1 (***B***), or solely injected with AS Cx38 co-injected in all oocytes to suppress endogenous Cx38 expression (***C***). Presence of standard NaCl bath solution with or without divalent cations (øMg^2+^øCa^2+^) or NMDG-Cl bath solution without divalent cations is indicated by black, red, and blue bars, respectively. Overlays of whole-cell current traces resulting from voltage step protocols are shown below the continuous current recordings. *Middle panels* Average I/V-plots (mean ± SEM) were constructed from similar recordings as shown in *left panels* (***A***: *n* =19, N=2; ***B***: *n* =18, N=2; ***C***: *n* =8, N=1). *Right panels* Summary data of the same experiments as shown in (***A***-***C***). The maximal inward currents reached during the application of hyperpolarizing pulses of-100 mV in three different bath solutions are shown. Lines connect data points obtained from one oocyte. The *p*-values were calculated by the Friedman test with Dunn’s post hoc test. ***D***, Maximal currents in PC2 *vs.* PC2/PC1 expressing oocytes. The maximal inward currents measured in NaCl bath solution without divalent cations (NaCl øMg^2+^øCa^2+^) at-100 mV are shown. Data are from the same experiments as summarized in the I/V plots shown in (***A***-***C***). The *p*-values were calculated by the Kruskal-Wallis test with Dunn’s post hoc test. ***E***, Western blot analysis of cell surface (*left panels*) and intracellular (*right panels*) expression of PC2 and PC1 in oocytes from one batch. Original uncropped images of the same blots are shown in Appendix Fig.S1.

**Fig. 2EV.**
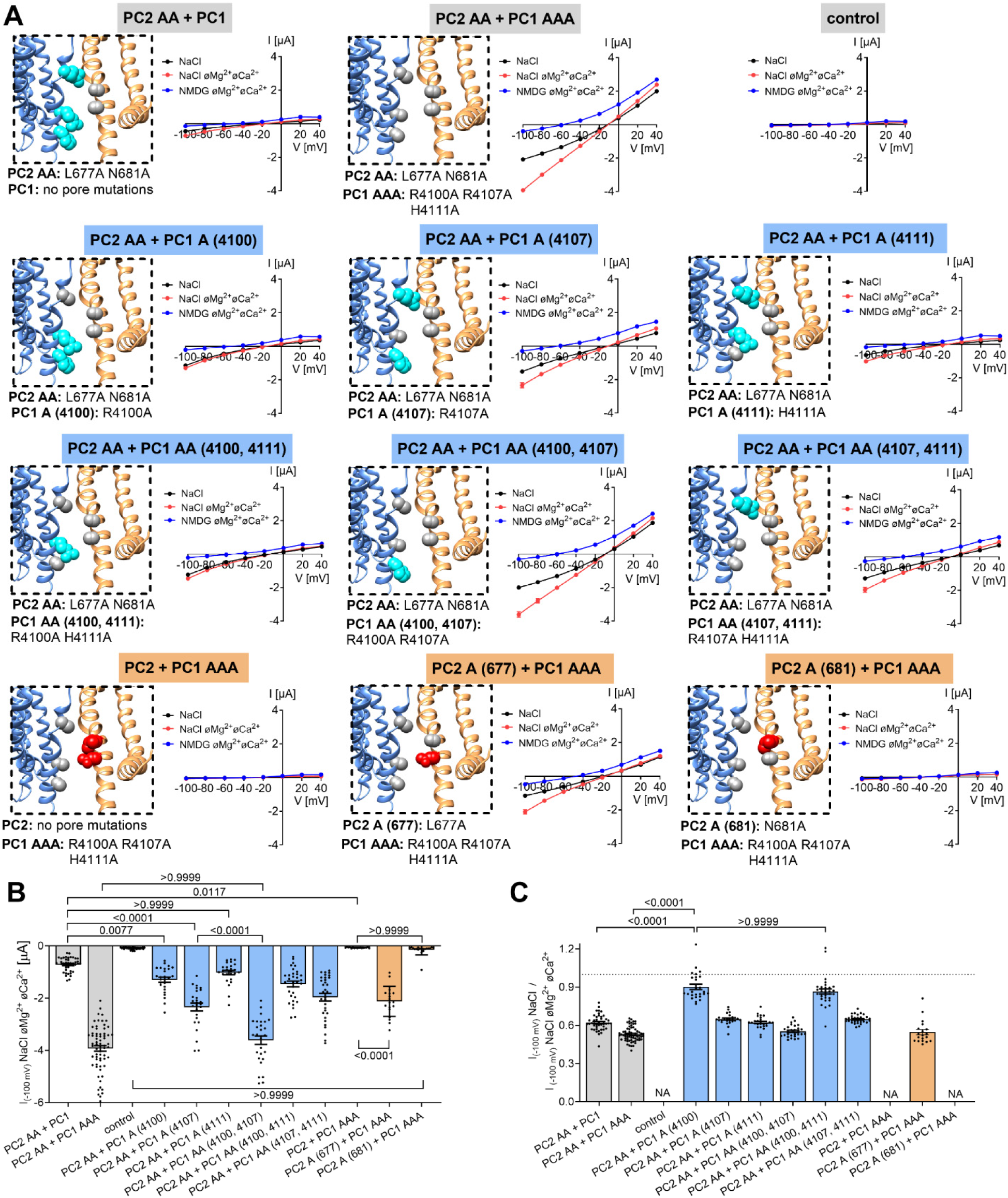
Alanine substitutions of the R4107 residue in PC1 and L677 residue in PC2 are the major contributors to the GOF effect. ***A****, Left panels* Localization of alanine substitutions in grey within the pore of heteromeric PC2/PC1 ion channels introduced to create PC2 and PC1 constructs as indicated. *Right panels* Average I/V-plots (mean ± SEM) obtained using the same experimental approach as shown in Fig. 1 from oocytes co-expressing the corresponding PC2 (2.5 ng) and PC1 (5 ng) constructs in three different bath solutions (PC2 AA + PC1: *n* =44, N=5; PC2 AA + PC1 AAA: *n* =66, N=7; control: *n* =60, N=7; PC2 AA + PC1 A (4100): *n* =26, N=3; PC2 AA + PC1 A (4107): *n* =26, N=3; PC2 AA + PC1 A (4111): *n* =26, N=3; PC2 AA + PC1 AA (4100, 4111): *n* =32, N=4; PC2 AA + PC1 AA (4100, 4107): *n* =28, N=4; PC2 AA + PC1 AA (4107, 4111): *n* =32, N=4; PC2 + PC1 AAA: *n* =17, N=2; PC2 A (677) + PC1 AAA: *n* =19, N=2; PC2 A (681) + PC1 AAA: *n* =19, N=2). ***B***, Maximal currents in oocytes expressing different PC2/PC1 heteromers. The maximal inward currents measured in NaCl bath solution without divalent cations (NaCl øMg^2+^øCa^2+^) at-100 mV are shown. Data are from the same experiments as in (***A***). The *p*-values were calculated by the one-way ANOVA with Bonferroni’s post hoc test. ***C***, Summary of the relative inhibitory effects of Ca^2+^ and Mg^2+^ on PC2/PC1-mediated sodium inward currents calculated as described in Fig. 1G. The *p*-values were calculated by the one-way ANOVA with Bonferroni’s post hoc test. NA, not applicable.

**Fig. 3EV.**
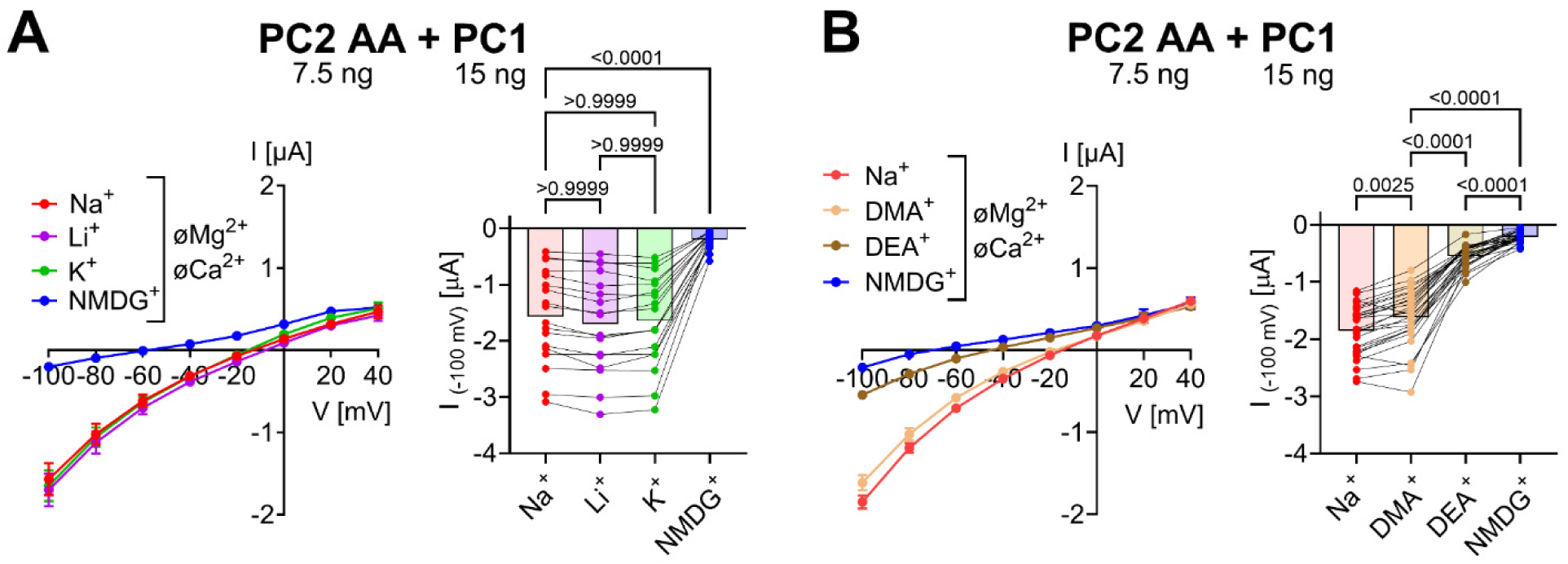
Permeability for monovalent cations of heteromeric PC2 AA / PC1 ion channels. ***A*, *B***, Permeability for small inorganic monovalent cations (***A***) and mid-size organic monovalent cations (***B***) was assessed as described in Fig. 2. *Left panels* Average I/V-plots (mean ± SEM). *Right panels* The maximal inward currents at-100 mV. The current values were corrected for endogenous oocyte currents shown in Appendix Fig. S8. Average values and individual data points are shown (***A***, *n* =18, N=2; ***B***, *n* =29, N=3). Lines connect data points obtained from one oocyte. The *p*-values were calculated by the repeated measures one-way ANOVA with Bonferroni’s post hoc test.

**Fig. 4EV.**
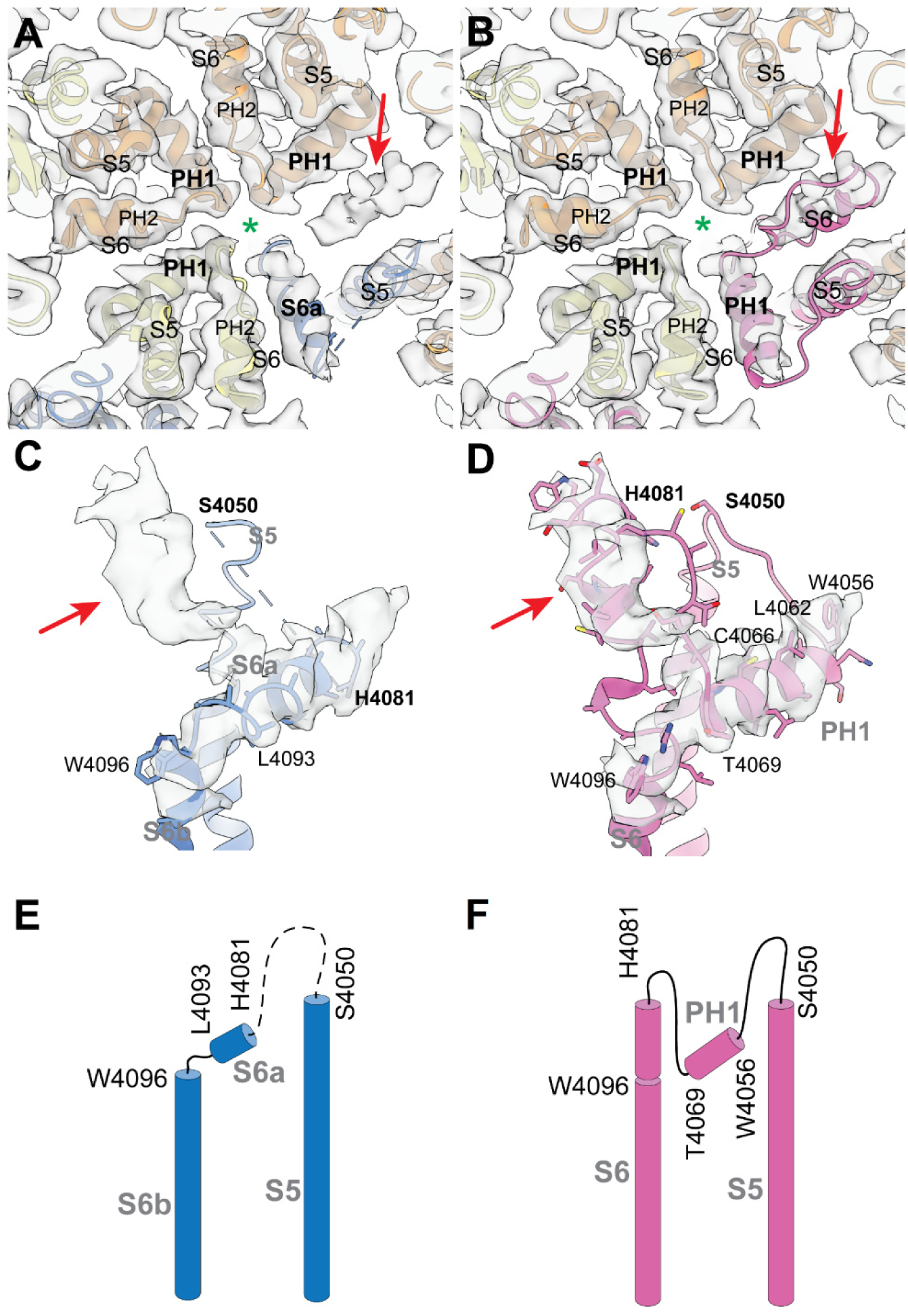
Re-interpretation of the published cryo-EM data. ***A, B*** View along the central cavity (indicated by the green color asterisk) on the selectivity filter region of the original model (***A***, PDB ID 6A70) or the re-interpreted model (***B***) of the PC2/PC1 heterotetrameric complex. PC2 subunits are colored as in Fig.1B, PC1 is in blue in ***A*** and in pink in ***B***. The respective EM density (EMD-6991) is shown as transparent volume. The red arrow indicates an unassigned, cylindrical density. ***C, D,*** Side view on the structural elements S5-S6a-S6b of the original model of PC1 (***C***) or on the S5-PH1-S6 elements of the re-interpreted model of PC1 (***D***). The respective PC1 EM density is shown as transparent volume. Several residues are labeled for orientation. Not modeled residues in ***C*** are indicated by a broken line. Side chains are shown in stick representation (for S6a in ***C*** side chains are not present according to the original interpretation). The red arrow points to the same unassigned density as indicated in ***A*** and ***B***. ***E, F*** Secondary structure diagrams of the original (***E***) and the re-interpreted model (***F***) of PC1.

**Fig. 5EV.**
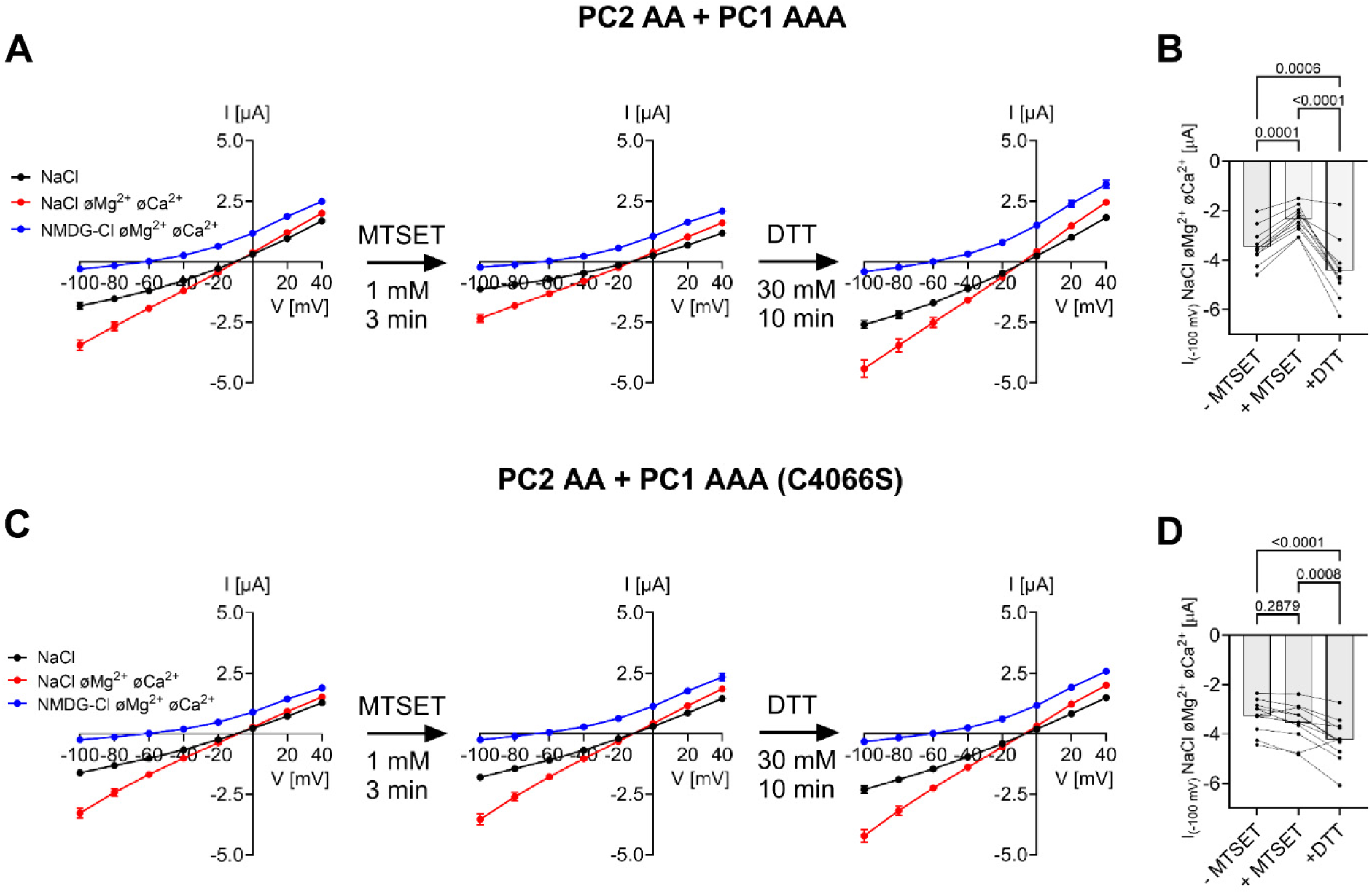
The inhibitory effect of MTSET on ion channel function of PC2 AA / PC1 AAA heteromers can be rescued by DTT. Average I/V-plots (mean ± SEM; ***A*, *C***) and the corresponding summary data showing the maximal inward current reached at-100 mV in divalent free NaCl (NaCl øMg^2+^ øCa^2+^) bath solution (mean values and individual data points; ***B*, *D***) are presented. In each individual oocyte the currents were measured before the treatment with MTSET (-MTSET), after 5 min incubation in NaCl bath solution supplemented with 1 mM of MTSET (+ MTSET), and after 10 min incubation in NaCl bath solution supplemented with 30 mM of DTT (+ DTT). The experimental protocol was similar to that described for Fig.4. Lines connect data points obtained from one oocyte. It is noteworthy, that the slow spontaneous current run-up observed in oocytes co-expressing PC2 AA and PC1 AAA with the C4066S mutation (***C*, *D***) was clearly different from the acute inhibitory response to MTSET and the subsequent rescue effect of DTT observed in oocytes co-expressing PC2 AA and PC1 AAA (***A*, *B***). The *p*-values were calculated by the repeated measures one-way ANOVA with Bonferroni’s post hoc test (*n* =11, N=2).

## Notes

### Competing Interest Statement

The authors have declared no competing interest.

## References

Anyatonwu GI, Estrada M, Tian X, Somlo S, Ehrlich BE (2007) Regulation of ryanodine receptor-dependent calcium signaling by polycystin-2. Proc Natl Acad Sci U S A 104: 6454–6459

Arif Pavel M, Lv C, Ng C, Yang L, Kashyap P, Lam C, Valentino V, Fung HY, Campbell T, Møller SG et al (2016) Function and regulation of TRPP2 ion channel revealed by a gain-of-function mutant. Proc Natl Acad Sci U S A 113: E2363–2372

Bergmann C, Guay-Woodford LM, Harris PC, Horie S, Peters DJM, Torres VE (2018) Polycystic kidney disease. Nat Rev Dis Primers 4: 50

Bjarnadóttir TK, Fredriksson R, Schiöth HB (2007) The adhesion GPCRs: a unique family of G protein-coupled receptors with important roles in both central and peripheral tissues. Cell Mol Life Sci 64: 2104–2119

Cai Y, Maeda Y, Cedzich A, Torres VE, Wu G, Hayashi T, Mochizuki T, Park JH, Witzgall R, Somlo S (1999) Identification and characterization of polycystin-2, the PKD2 gene product. J Biol Chem 274: 28557–28565

Cantero MDR, Cantiello HF (2022) Polycystin-2 (TRPP2): Ion channel properties and regulation. Gene 827: 146313

Cao E (2020) Structural mechanisms of transient receptor potential ion channels. J Gen Physiol 152(3): e201811998

Caplan MJ (2019) Holding open the door reveals a new view of polycystin channel function. EMBO Rep 20: e49156

Chauvet V, Qian F, Boute N, Cai Y, Phakdeekitacharoen B, Onuchic LF, Attié-Bitach T, Guicharnaud L, Devuyst O, Germino GG et al (2002) Expression of PKD1 and PKD2 transcripts and proteins in human embryo and during normal kidney development. Am J Pathol 160: 973–983

Chebib FT, Torres VE (2016) Autosomal dominant polycystic kidney disease: core curriculum 2016. Am J Kidney Dis 67: 792–810

Cornec-Le Gall E, Torres VE, Harris PC (2018) Genetic complexity of autosomal dominant polycystic kidney and liver diseases. J Am Soc Nephrol 29: 13–23

Douguet D, Patel A, Honoré E (2019) Structure and function of polycystins: insights into polycystic kidney disease. Nat Rev Nephrol 15: 412–422

Emsley P, Lohkamp B, Scott WG, Cowtan K (2010) Features and development of Coot. Acta Crystallogr D Biol Crystallogr 66: 486–501

Geng L, Segal Y, Peissel B, Deng N, Pei Y, Carone F, Rennke HG, Glücksmann-Kuis AM, Schneider MC, Ericsson M et al (1996) Identification and localization of polycystin, the PKD1 gene product. J Clin Invest 98: 2674–2682

Grantham JJ, Mulamalla S, Swenson-Fields KI (2011) Why kidneys fail in autosomal dominant polycystic kidney disease. Nat Rev Nephrol 7: 556–566

Grieben M, Pike AC, Shintre CA, Venturi E, El-Ajouz S, Tessitore A, Shrestha L, Mukhopadhyay S, Mahajan P, Chalk R et al (2017) Structure of the polycystic kidney disease TRP channel Polycystin-2 (PC2). Nat Struct Mol Biol 24: 114–122

Grosch M, Brunner K, Ilyaskin AV, Schober M, Staudner T, Schmied D, Stumpp T, Schmidt KN, Madej MG, Pessoa TD et al (2021) A polycystin-2 protein with modified channel properties leads to an increased diameter of renal tubules and to renal cysts. J Cell Sci 134(16): jcs259013

Ha K, Mundt-Machado N, Bisignano P, Pinedo A, Raleigh DR, Loeb G, Reiter JF, Cao E, Delling M (2024) Cilia-enriched oxysterol 7β,27-DHC is required for polycystin ion channel activation. Nat Commun 15(1): 6468

Ha K, Nobuhara M, Wang Q, Walker RV, Qian F, Schartner C, Cao E, Delling M (2020) The heteromeric PC-1/PC-2 polycystin complex is activated by the PC-1 N-terminus. Elife 9: e60684

Hanaoka K, Qian F, Boletta A, Bhunia AK, Piontek K, Tsiokas L, Sukhatme VP, Guggino WB, Germino GG (2000) Co-assembly of polycystin-1 and-2 produces unique cation-permeable currents. Nature 408: 990–994

Hille B (2001) Ion Channels of Excitable Membranes, 3rd Ed., Sinauer Associates Inc, Sunderland

Hoffmeister H, Babinger K, Gürster S, Cedzich A, Meese C, Schadendorf K, Osten L, de Vries U, Rascle A, Witzgall R (2011) Polycystin-2 takes different routes to the somatic and ciliary plasma membrane. J Cell Biol 192: 631–645

Hughes J, Ward CJ, Peral B, Aspinwall R, Clark K, San Millán JL, Gamble V, Harris PC (1995) The polycystic kidney disease 1 (PKD1) gene encodes a novel protein with multiple cell recognition domains. Nat Genet 10: 151–160

Hulse RE, Li Z, Huang RK, Zhang J, Clapham DE (2018) Cryo-EM structure of the polycystin 2-l1 ion channel. Elife 7: e36931

Ilyaskin AV, Korbmacher C, Diakov A (2021) Inhibition of the epithelial sodium channel (ENaC) by connexin 30 involves stimulation of clathrin-mediated endocytosis. J Biol Chem 296: 100404

Kellenberger S, Gautschi I, Schild L (2002) An external site controls closing of the epithelial Na^+^ channel ENaC. J Physiol 543: 413–424

Kleene SJ, Kleene NK (2017) The native TRPP2-dependent channel of murine renal primary cilia. Am J Physiol Renal Physiol 312: F96–108

Koptides M, Hadjimichael C, Koupepidou P, Pierides A, Constantinou Deltas C (1999) Germinal and somatic mutations in the PKD2 gene of renal cysts in autosomal dominant polycystic kidney disease. Hum Mol Genet 8: 509–513

Koulen P, Cai Y, Geng L, Maeda Y, Nishimura S, Witzgall R, Ehrlich BE, Somlo S (2002) Polycystin-2 is an intracellular calcium release channel. Nat Cell Biol 4: 191–197

Li Y, Wright JM, Qian F, Germino GG, Guggino WB (2005) Polycystin 2 interacts with type I inositol 1,4,5-trisphosphate receptor to modulate intracellular Ca^2+^ signaling. J Biol Chem 280: 41298–41306

Liu X, Vien T, Duan J, Sheu SH, DeCaen PG, Clapham DE (2018) Polycystin-2 is an essential ion channel subunit in the primary cilium of the renal collecting duct epithelium. Elife 7: e33183

Lorenz C, Pusch M, Jentsch TJ (1996) Heteromultimeric CLC chloride channels with novel properties. Proc Natl Acad Sci U S A 93: 13362–13366

Ma M, Gallagher AR, Somlo S (2017) Ciliary mechanisms of cyst formation in polycystic kidney disease. Cold Spring Harb Perspect Biol 9(11): a028209

Ma M, Tian X, Igarashi P, Pazour GJ, Somlo S (2013) Loss of cilia suppresses cyst growth in genetic models of autosomal dominant polycystic kidney disease. Nat Genet 45: 1004–1012

Markowitz GS, Cai Y, Li L, Wu G, Ward LC, Somlo S, D’Agati VD (1999) Polycystin-2 expression is developmentally regulated. Am J Physiol 277: F17–25

Mochizuki T, Wu G, Hayashi T, Xenophontos SL, Veldhuisen B, Saris JJ, Reynolds DM, Cai Y, Gabow PA, Pierides A et al (1996) PKD2, a gene for polycystic kidney disease that encodes an integral membrane protein. Science 272: 1339–1342

Murshudov GN, Vagin AA, Dodson EJ (1997) Refinement of macromolecular structures by the maximum-likelihood method. Acta Crystallogr D Biol Crystallogr 53: 240–255

Nieberler M, Kittel RJ, Petrenko AG, Lin HH, Langenhan T (2016) Control of adhesion GPCR function through proteolytic processing. Handb Exp Pharmacol 234: 83–109

Nims N, Vassmer D, Maser RL (2003) Transmembrane domain analysis of polycystin-1, the product of the polycystic kidney disease-1 (PKD1) gene: evidence for 11 membrane-spanning domains. Biochemistry 42: 13035–13048

Ong AC, Ward CJ, Butler RJ, Biddolph S, Bowker C, Torra R, Pei Y, Harris PC (1999) Coordinate expression of the autosomal dominant polycystic kidney disease proteins, polycystin-2 and polycystin-1, in normal and cystic tissue. Am J Pathol 154: 1721–1729

Padhy B, Xie J, Wang R, Lin F, Huang CL (2022) Channel function of polycystin-2 in the endoplasmic reticulum protects against autosomal dominant polycystic kidney disease. J Am Soc Nephrol 33: 1501–1516

Parnell SC, Magenheimer BS, Maser RL, Pavlov TS, Havens MA, Hastings ML, Jackson SF, Ward CJ, Peterson KR, Staruschenko A et al (2018) A mutation affecting polycystin-1 mediated heterotrimeric G-protein signaling causes PKD. Hum Mol Genet 27: 3313–3324

Parnell SC, Magenheimer BS, Maser RL, Rankin CA, Smine A, Okamoto T, Calvet JP (1998) The polycystic kidney disease-1 protein, polycystin-1, binds and activates heterotrimeric G-proteins in vitro. Biochem Biophys Res Commun 251: 625–631

Parnell SC, Magenheimer BS, Maser RL, Zien CA, Frischauf AM, Calvet JP (2002) Polycystin-1 activation of c-Jun N-terminal kinase and AP-1 is mediated by heterotrimeric G proteins. J Biol Chem 277: 19566–19572

Pawnikar S, Magenheimer BS, Joshi K, Nevarez-Munoz E, Haldane A, Maser RL, Miao Y (2024) Activation of polycystin-1 signaling by binding of stalk-derived peptide agonists. Elife 13: RP95992

Pawnikar S, Magenheimer BS, Munoz EN, Maser RL, Miao Y (2022) Mechanism of tethered agonist-mediated signaling by polycystin-1. Proc Natl Acad Sci U S A 119: e2113786119

Pazour GJ, San Agustin JT, Follit JA, Rosenbaum JL, Witman GB (2002) Polycystin-2 localizes to kidney cilia and the ciliary level is elevated in orpk mice with polycystic kidney disease. Curr Biol 12: R378–380

Pei Y, Watnick T, He N, Wang K, Liang Y, Parfrey P, Germino G, St George-Hyslop P (1999) Somatic PKD2 mutations in individual kidney and liver cysts support a “two-hit” model of cystogenesis in type 2 autosomal dominant polycystic kidney disease. J Am Soc Nephrol 10: 1524–1529

Pettersen EF, Goddard TD, Huang CC, Couch GS, Greenblatt DM, Meng EC, Ferrin TE (2004) UCSF Chimera--a visualization system for exploratory research and analysis. J Comput Chem 25: 1605–1612

Qian F, Boletta A, Bhunia AK, Xu H, Liu L, Ahrabi AK, Watnick TJ, Zhou F, Germino GG (2002) Cleavage of polycystin-1 requires the receptor for egg jelly domain and is disrupted by human autosomal-dominant polycystic kidney disease 1-associated mutations. Proc Natl Acad Sci U S A 99: 16981–16986

Qian F, Germino FJ, Cai Y, Zhang X, Somlo S, Germino GG (1997) PKD1 interacts with PKD2 through a probable coiled-coil domain. Nat Genet 16: 179–183

Qian F, Watnick TJ, Onuchic LF, Germino GG (1996) The molecular basis of focal cyst formation in human autosomal dominant polycystic kidney disease type I. Cell 87: 979–987

Sali A, Blundell TL (1993) Comparative protein modelling by satisfaction of spatial restraints. J Mol Biol 234: 779–815

Sammels E, Devogelaere B, Mekahli D, Bultynck G, Missiaen L, Parys JB, Cai Y, Somlo S, De Smedt H (2010) Polycystin-2 activation by inositol 1,4,5-trisphosphate-induced Ca^2+^ release requires its direct association with the inositol 1,4,5-trisphosphate receptor in a signaling microdomain. J Biol Chem 285: 18794–18805

Shen PS, Yang X, DeCaen PG, Liu X, Bulkley D, Clapham DE, Cao E (2016) The structure of the polycystic kidney disease channel PKD2 in lipid nanodiscs. Cell 167(3): 763–773.e11

Staudner T, Geiges L, Khamseekaew J, Sure F, Korbmacher C, Ilyaskin AV (2024) Disease-associated missense mutations in the pore loop of polycystin-2 alter its ion channel function in a heterologous expression system. J Biol Chem 300(8): 107574

Su Q, Chen M, Wang Y, Li B, Jing D, Zhan X, Yu Y, Shi Y (2021) Structural basis for Ca^2+^ activation of the heteromeric PKD1L3/PKD2L1 channel. Nat Commun 12(1): 4871

Su Q, Hu F, Ge X, Lei J, Yu S, Wang T, Zhou Q, Mei C, Shi Y (2018a) Structure of the human PKD1-PKD2 complex. Science 361(6406): eaat9819

Su Q, Hu F, Liu Y, Ge X, Mei C, Yu S, Shen A, Zhou Q, Yan C, Lei J et al (2018b) Cryo-EM structure of the polycystic kidney disease-like channel PKD2L1. Nat Commun 9(1): 1192

Sure F, Bertog M, Afonso S, Diakov A, Rinke R, Madej MG, Wittmann S, Gramberg T, Korbmacher C, Ilyaskin AV (2022) Transmembrane serine protease 2 (TMPRSS2) proteolytically activates the epithelial sodium channel (ENaC) by cleaving the channel’s γ-subunit. J Biol Chem 298(6): 102004

Tan AY, Zhang T, Michaeel A, Blumenfeld J, Liu G, Zhang W, Zhang Z, Zhu Y, Rennert L, Martin C et al (2018) Somatic mutations in renal cyst epithelium in autosomal dominant polycystic kidney disease. J Am Soc Nephrol 29: 2139–2156

Tsiokas L, Kim E, Arnould T, Sukhatme VP, Walz G (1997) Homo-and heterodimeric interactions between the gene products of PKD1 and PKD2. Proc Natl Acad Sci U S A 94: 6965–6970

Vien TN, Wang J, Ng LCT, Cao E, DeCaen PG (2020) Molecular dysregulation of ciliary polycystin-2 channels caused by variants in the TOP domain. Proc Natl Acad Sci U S A 117: 10329–10338

Walker RV, Keynton JL, Grimes DT, Sreekumar V, Williams DJ, Esapa C, Wu D, Knight MM, Norris DP (2019) Ciliary exclusion of polycystin-2 promotes kidney cystogenesis in an autosomal dominant polycystic kidney disease model. Nat Commun 10(1): 4072

Wang Q, Corey RA, Hedger G, Aryal P, Grieben M, Nasrallah C, Baronina A, Pike ACW, Shi J, Carpenter EP, et al Cryo-EM structure of human polycystin-2/PKD2 in UDM supplemented with PI(4,5)P2 (2020a) 6T9N [DATASET]

Wang Q, Corey RA, Hedger G, Aryal P, Grieben M, Nasrallah C, Baronina A, Pike ACW, Shi J, Carpenter EP et al (2020b) Lipid interactions of a ciliary membrane TRP channel: simulation and structural studies of polycystin-2. Structure 28: 169–184.e165

Wang Y, Wang Z, Pavel MA, Ng C, Kashyap P, Li B, Morais TDC, Ulloa GA, Yu Y (2023) The diverse effects of pathogenic point mutations on ion channel activity of a gain-of-function polycystin-2. J Biol Chem 299(5): 104674

Wang Z, Chen M, Su Q, Morais TDC, Wang Y, Nazginov E, Pillai AR, Qian F, Shi Y, Yu Y (2024) Molecular and structural basis of the dual regulation of the polycystin-2 ion channel by small-molecule ligands. Proc Natl Acad Sci U S A 121: e2316230121

Wang Z, Ng C, Liu X, Wang Y, Li B, Kashyap P, Chaudhry HA, Castro A, Kalontar EM, Ilyayev L et al (2019) The ion channel function of polycystin-1 in the polycystin-1/polycystin-2 complex. EMBO Rep 20: e48336

Wegierski T, Steffl D, Kopp C, Tauber R, Buchholz B, Nitschke R, Kuehn EW, Walz G, Köttgen M (2009) TRPP2 channels regulate apoptosis through the Ca^2+^ concentration in the endoplasmic reticulum. EMBO J 28: 490–499

Wei W, Hackmann K, Xu H, Germino G, Qian F (2007) Characterization of cis-autoproteolysis of polycystin-1, the product of human polycystic kidney disease 1 gene. J Biol Chem 282: 21729–21737

Wilkes M, Madej MG, Kreuter L, Rhinow D, Heinz V, De Sanctis S, Ruppel S, Richter RM, Joos F, Grieben M et al (2017) Molecular insights into lipid-assisted Ca^2+^ regulation of the TRP channel Polycystin-2. Nat Struct Mol Biol 24: 123–130

Wu G, D’Agati V, Cai Y, Markowitz G, Park JH, Reynolds DM, Maeda Y, Le TC, Hou H, Jr., Kucherlapati R et al (1998) Somatic inactivation of Pkd2 results in polycystic kidney disease. Cell 93: 177–188

Wu LJ, Sweet TB, Clapham DE (2010) International Union of Basic and Clinical Pharmacology. LXXVI. Current progress in the mammalian TRP ion channel family. Pharmacol Rev 62: 381–404

Yoder BK, Hou X, Guay-Woodford LM (2002) The polycystic kidney disease proteins, polycystin-1, polycystin-2, polaris, and cystin, are co-localized in renal cilia. J Am Soc Nephrol 13: 2508–2516

Yu Y, Ulbrich MH, Li MH, Buraei Z, Chen XZ, Ong AC, Tong L, Isacoff EY, Yang J (2009) Structural and molecular basis of the assembly of the TRPP2/PKD1 complex. Proc Natl Acad Sci U S A 106: 11558–11563

Zheng W, Yang X, Hu R, Cai R, Hofmann L, Wang Z, Hu Q, Liu X, Bulkley D, Yu Y et al (2018) Hydrophobic pore gates regulate ion permeation in polycystic kidney disease 2 and 2L1 channels. Nat Commun 9(1): 2302

Zhu J, Yu Y, Ulbrich MH, Li MH, Isacoff EY, Honig B, Yang J (2011) Structural model of the TRPP2/PKD1 C-terminal coiled-coil complex produced by a combined computational and experimental approach. Proc Natl Acad Sci U S A 108: 10133–10138

