## Appendix for "Ion channel function of polycystin-2/polycystin-1 heteromer revealed by structure-guided mutagenesis"

**Appendix Table of contents**

Figure S1; Figure S2; Figure S3; Figure S4; Figure S5; Figure S6; Figure S7; Figure S8; Figure S9; Figure S10; Figure S11; Appendix Table S1; Figure S12.

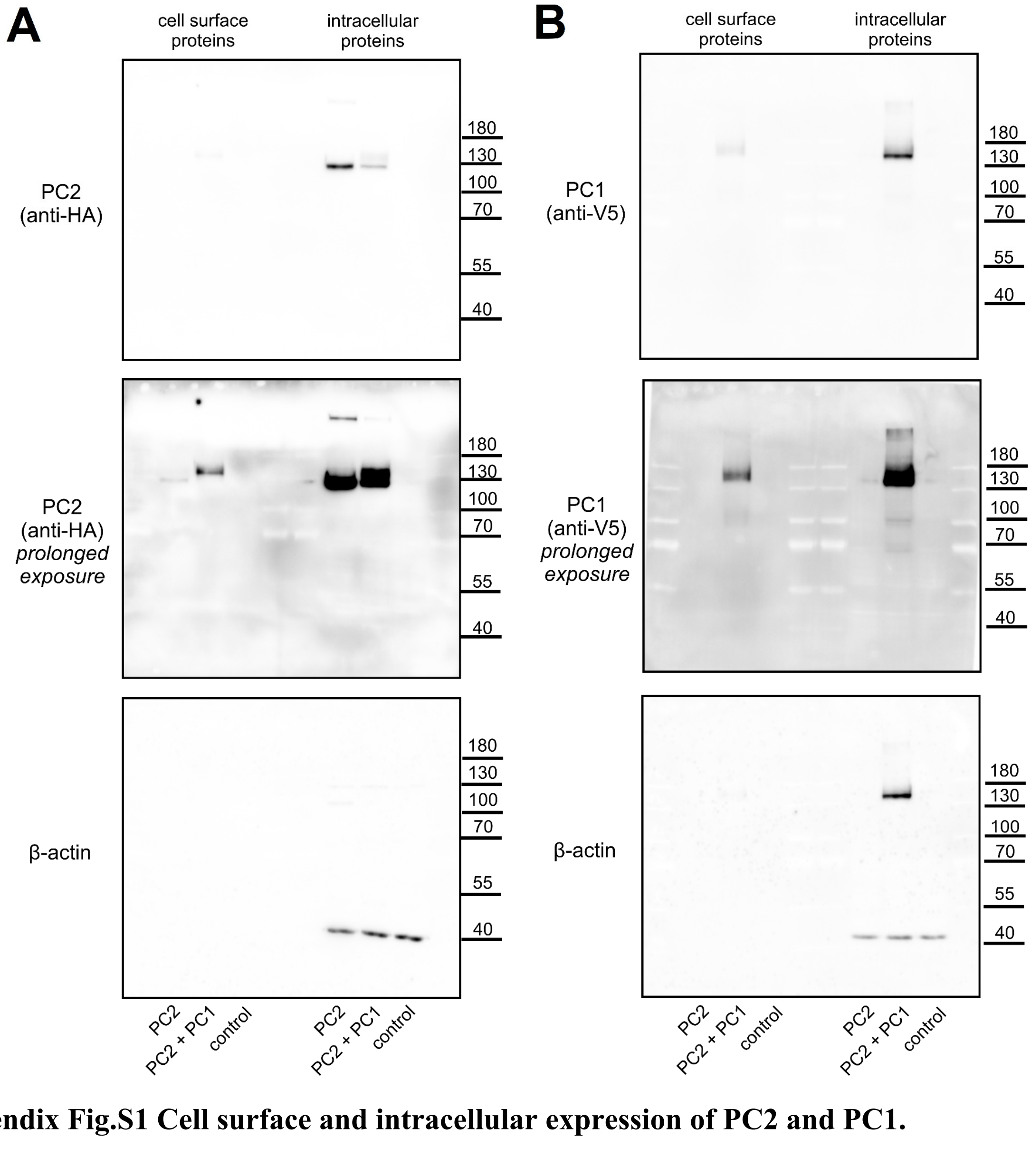


**Appendix Fig.S1 Cell surface and intracellular expression of PC2 and PC1.**

***A***, ***B*** Original uncropped images of the same western blots shown in Figure 1EV-***E***. Images showing PC2 (***A***, *upper* and *middle panels*) or PC1 (***B***, *upper* and *middle panels*) signals were obtained using short (*upper panels*) or prolonged (*middle panels*) exposure for optimal detection of intracellular or cell surface expression, respectively. To confirm separation of cell surface proteins from intracellular proteins, the membranes were stripped and re-probed using an anti-β-actin antibody (*lower panels*).

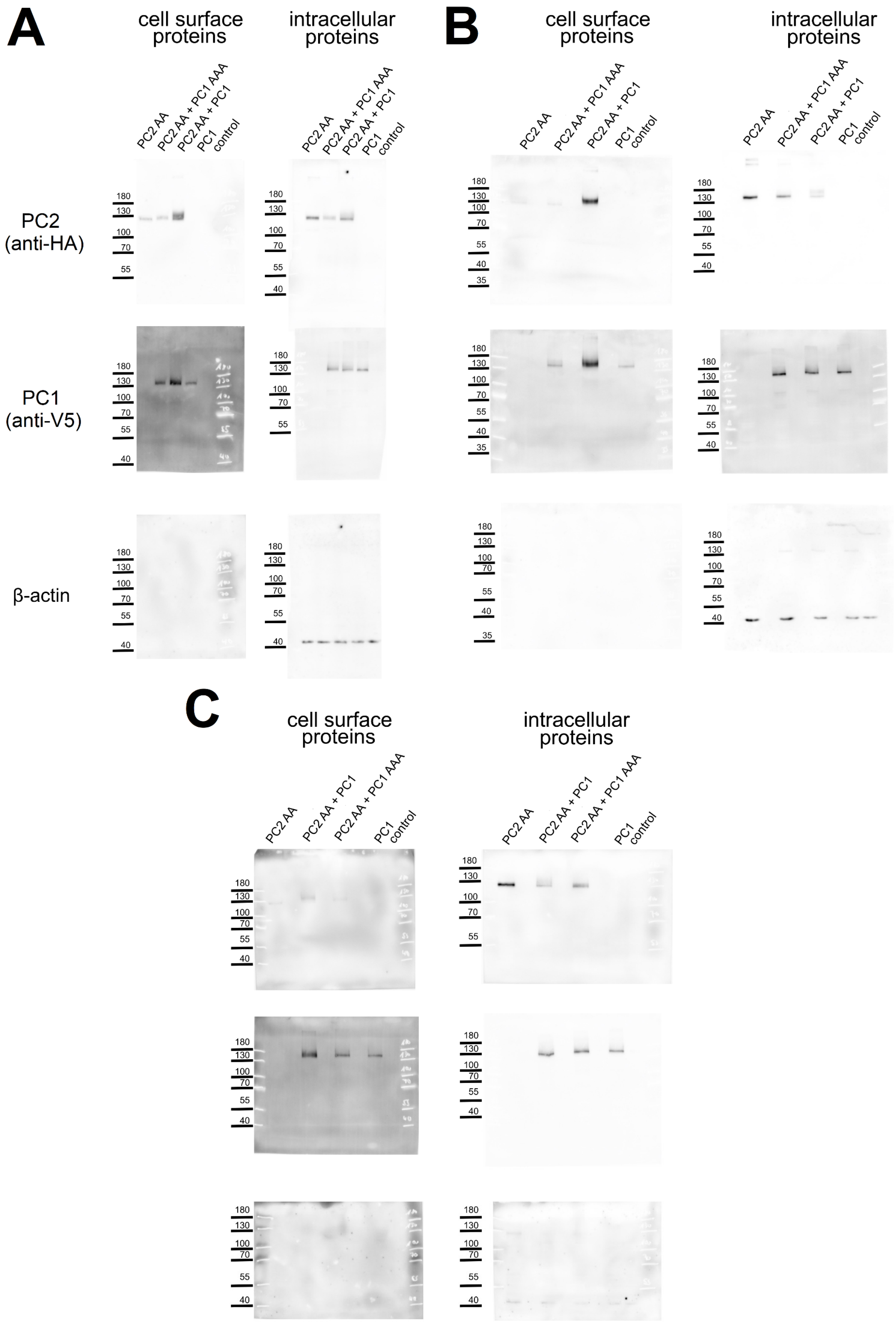


**Appendix Fig.S2 Cell surface and intracellular expression of PC2 AA, PC1 and PC1 AAA.**

Original uncropped western blot images obtained in oocytes from three different batches (***A***: batch 1; ***B***: batch 2; ***C***: batch 3). The data from the batch 1 (***A***) are included in Fig. 1 as panel ***H***.

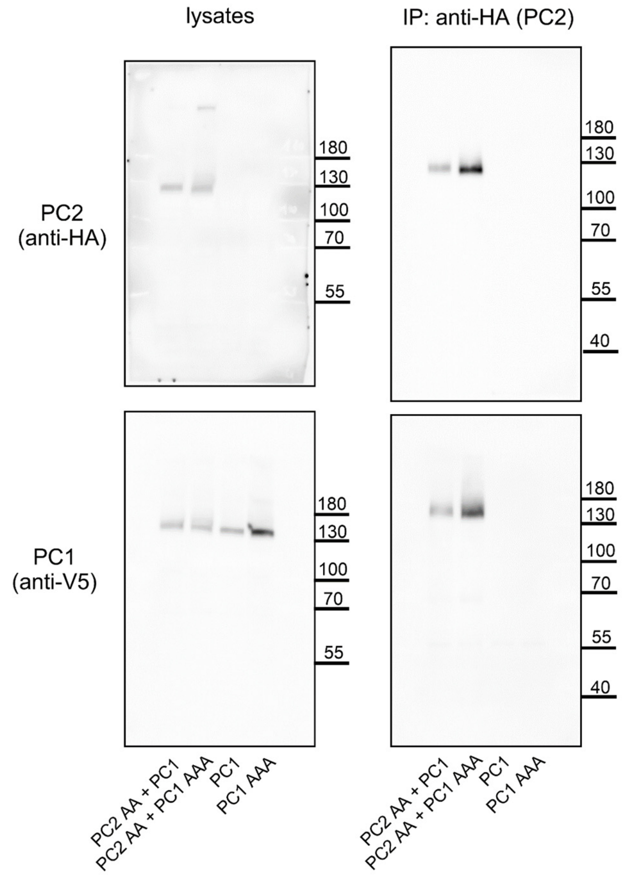


**Appendix Fig.S3 Validation of PC2/PC1 complex formation using co-IP and PC2-HA as a "bait" protein.**

Original uncropped images of the same western blots as shown in Figure 1-***I***.

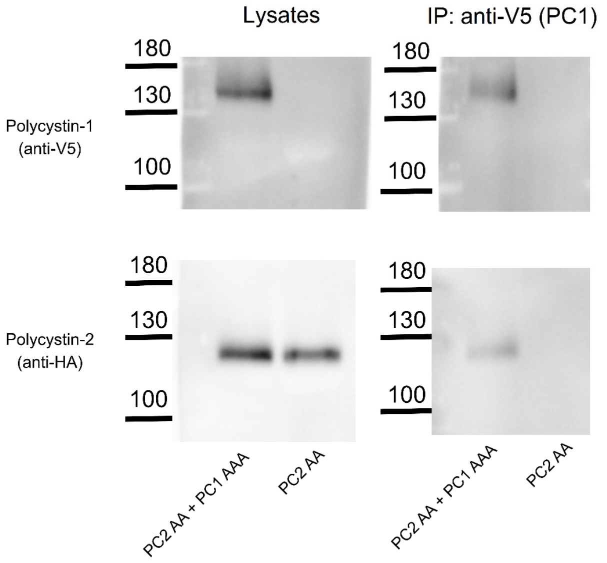


**Appendix Fig.S4 Validation of PC2/PC1 complex formation using co-IP and PC1-V5 as a "bait" protein.**

PC2 and PC1 were detected in co-IP preparations (*right panels*) and in corresponding cell lysates (*left panels*). PC2/PC1 complexes were isolated using an anti-V5 antibody conjugated to magnetic beads, which recognized V5-tagged PC1.

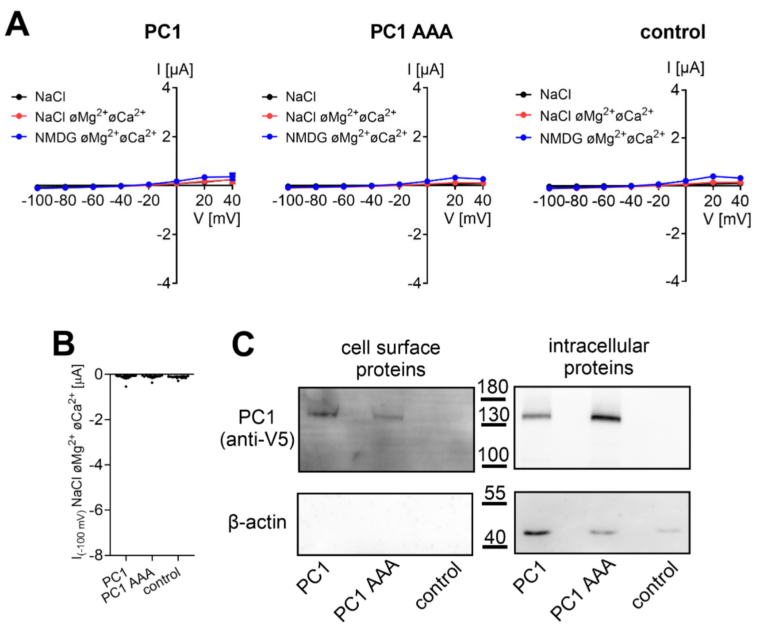


**Appendix Fig.S5 Expression of PC1 or PC1 AAA alone did not result in detectable ion channel currents.**

***A***, Average I/V-plots (mean ± SEM) obtained from oocytes injected with 15 ng of PC1 or PC1 AAA encoding cRNA or from control oocytes ($n$ =15, N=2) using the same experimental approach as shown in Fig. 1. ***B***, Maximal inward currents measured in NaCl bath solution without divalent cations (NaCl ø$Mg^{2+}$ø$Ca^{2+}$) at -100 mV. Data are from the same experiments as in (***A***). ***C***, Western blot analysis of cell surface (*left panels*) and intracellular (*right panels*) expression of PC1 and PC1 AAA in oocytes from one batch.

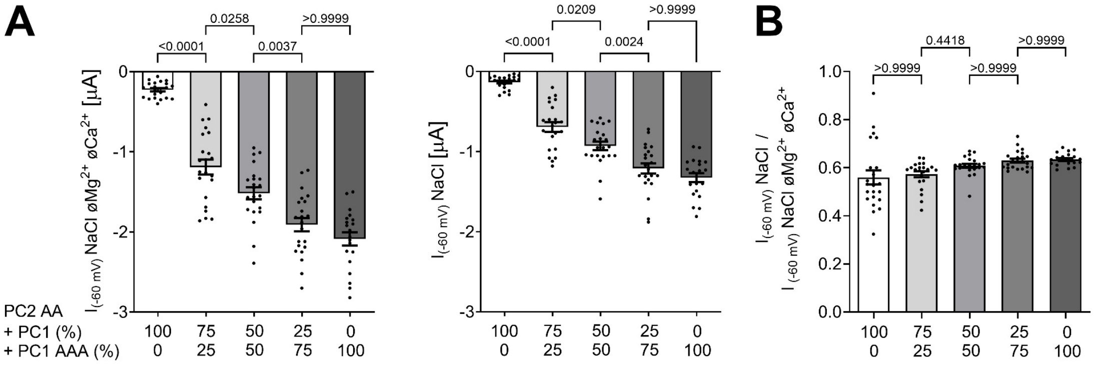


**Appendix Fig.S6 Replacing PC1 by PC1 AAA in co-expression experiments with PC2 AA increased ion channel currents without changing the inhibitory effect of divalent cations.**

***A*,** Inward currents were obtained in NaCl bath solution without (*left panel*) or with divalent cations (NaCl, *right panel*) from oocytes co-injected with a constant amount of PC2 AA cRNA (2.5 ng) and variable relative amounts of PC1 or PC1 AAA cRNAs (mean ± SEM; 20≤$n$≤22, N=2). The total cRNA amount of PC1+PC1AAA was kept constant at 5 ng. The relative amounts of PC1 and PC1 AAA cRNAs (in %) are indicated on the x-axis. The $p$-values were calculated by the one-way ANOVA with Bonferroni's post hoc test. ***B***, Summary of the relative inhibitory effects of $Ca^{2+}$ and $Mg^{2+}$ on PC2/PC1-mediated sodium inward currents. The $p$-values were calculated by the Kruskal-Wallis test with Dunn's post hoc test.

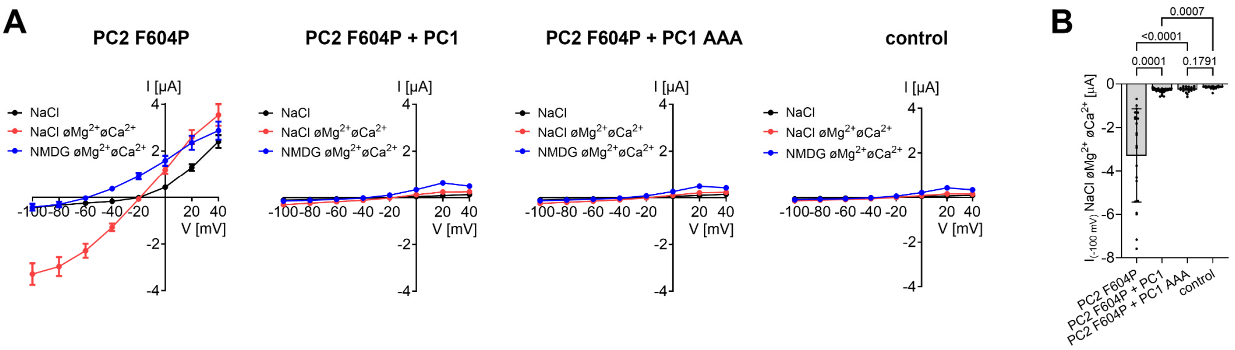


**Appendix Fig.S7 Ion channel function of PC2 F604P is blocked by PC1 or PC1 AAA co-expression.**

***A***, Average I/V-plots (mean ± SEM) obtained using the same experimental approach as shown in Fig. 1 from oocytes injected with 7.5 ng of cRNA encoding PC2 F604P only, co-injected with 15 ng of cRNA encoding PC1 or PC1 AAA, or from control oocytes (PC2 F604P: *n* =22, N=3; PC2 F604P + PC1: *n* =26, N=3; PC2 F604P + PC1 AAA: *n* =20, N=3; control: *n* =23, N=3). ***B***, Maximal inward currents measured in NaCl bath solution without divalent cations (NaCl øMg$^{2+}$øCa$^{2+}$) at -100 mV. Data are from the same experiments as in (***A***). The *p*-values were calculated by the Kruskal-Wallis test with Dunn's post hoc test.

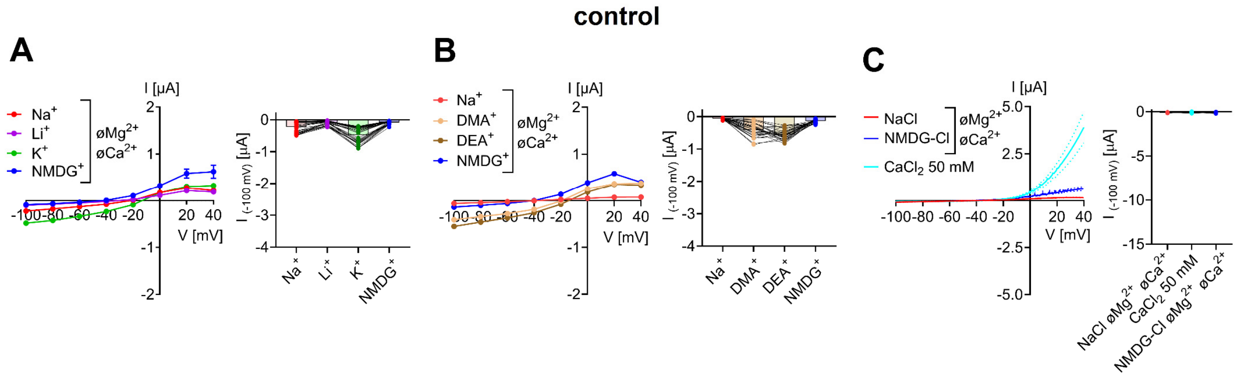


**Appendix Fig.S8 Effect of monovalent cation substitutions and application of 50 mM $CaCl_2$ bath solution on baseline currents in control oocytes.**

***A***-***C***, Baseline currents measured in control oocytes injected solely with AS Cx38 using a similar experimental protocol as described is Fig.2. *Left panels* Average I/V-plots (mean ± SEM). *Right panels* The maximal inward currents at -100 mV. Average values and individual data points are shown (***A***, *n* =36, N=3; ***B***, *n* =28, N=3; ***C***, *n* =21, N=3). Lines connect data points obtained from one oocyte. In Fig. 2 and Fig.3EV, these average whole-cell currents in different bath solutions were used to correct corresponding current values obtained in oocytes expressing polycystin constructs for endogenous oocyte currents.

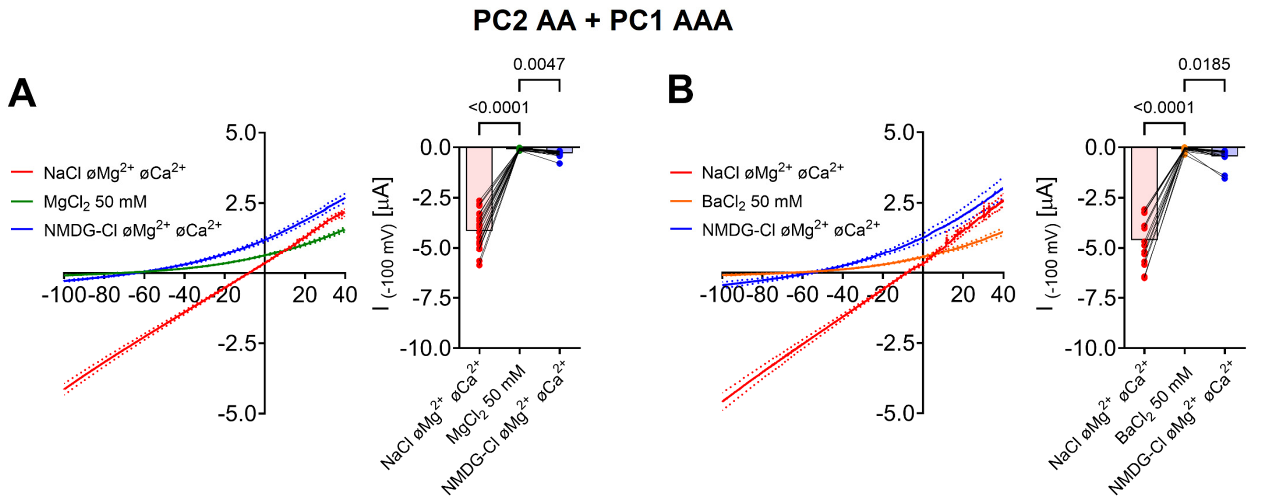


**Appendix Fig.S9 Heteromeric PC2 AA / PC1 AAA ion channels are impermeable for divalent cations $Mg^{2+}$ and $Ba^{2+}$.**

***A***, ***B*** Permeability for $Mg^{2+}$ (***A***) or $Ba^{2+}$ (***B***) was estimated using a similar experimental protocol as described is Fig.2.***E***, ***F*** but with 50 mM $MgCl_2$ or 50 mM $BaCl_2$ bath solutions, respectively. *Left panels* Average I/V-plots (mean ± SEM). *Right panels* Maximal inward currents at -100 mV. Average values and individual data points are shown (***A***, *n* =20, N=2; ***B***, *n* =15, N=2). Lines connect data points obtained from one oocyte. The *p*-values were calculated by the Friedman test with Dunn's post hoc test.

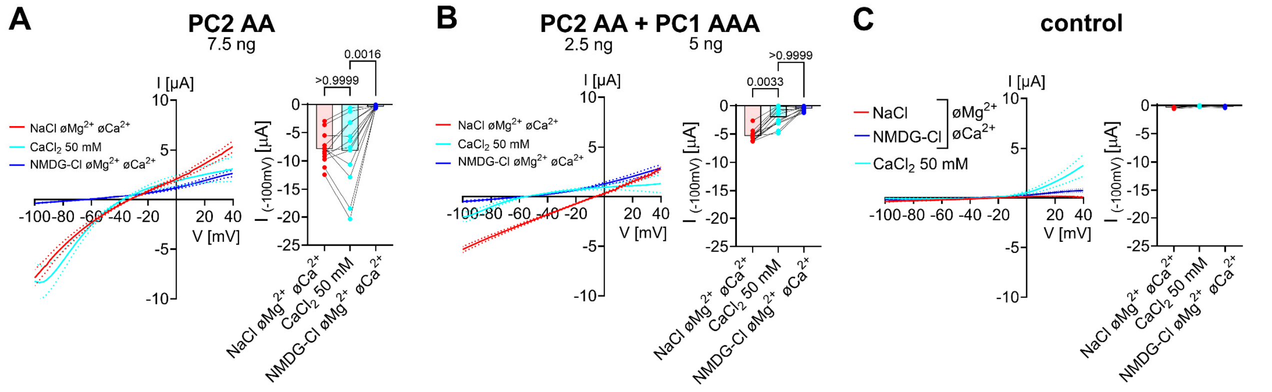


**Appendix Fig.S10 Heteromeric PC2 AA / PC1 AAA ion channels are less permeable for $Ca^{2+}$ than PC2 AA homomers even under experimental conditions facilitating $Ca^{2+}$ entry.**

***A***-***C*** Permeability for $Ca^{2+}$ in oocytes expressing PC2 AA (***A***), co-expressing PC2 AA and PC1 AAA (***B***), and control oocytes (***C***) was estimated using a similar experimental protocol as described is Fig.2.***E***, ***F*** but with a continuous holding potential of 0 mV instead of -60 mV between ramp protocols to reduce the voltage-dependent pore blocking effect of $Ca^{2+}$ (see Methods). The current values shown in ***A*** and ***B*** were corrected for endogenous oocyte currents shown in ***C***. *Left panels* Average I/V-plots (mean ± SEM). *Right panels* Maximal inward currents at -100 mV. Average values and individual data points are shown (***A***, *n* =12, N=2; ***B***, *n* =12, N=2; ***C***, *n* =6, N=2). Lines connect data points obtained from one oocyte. The *p*-values were calculated by the Friedman test with Dunn's post hoc test.

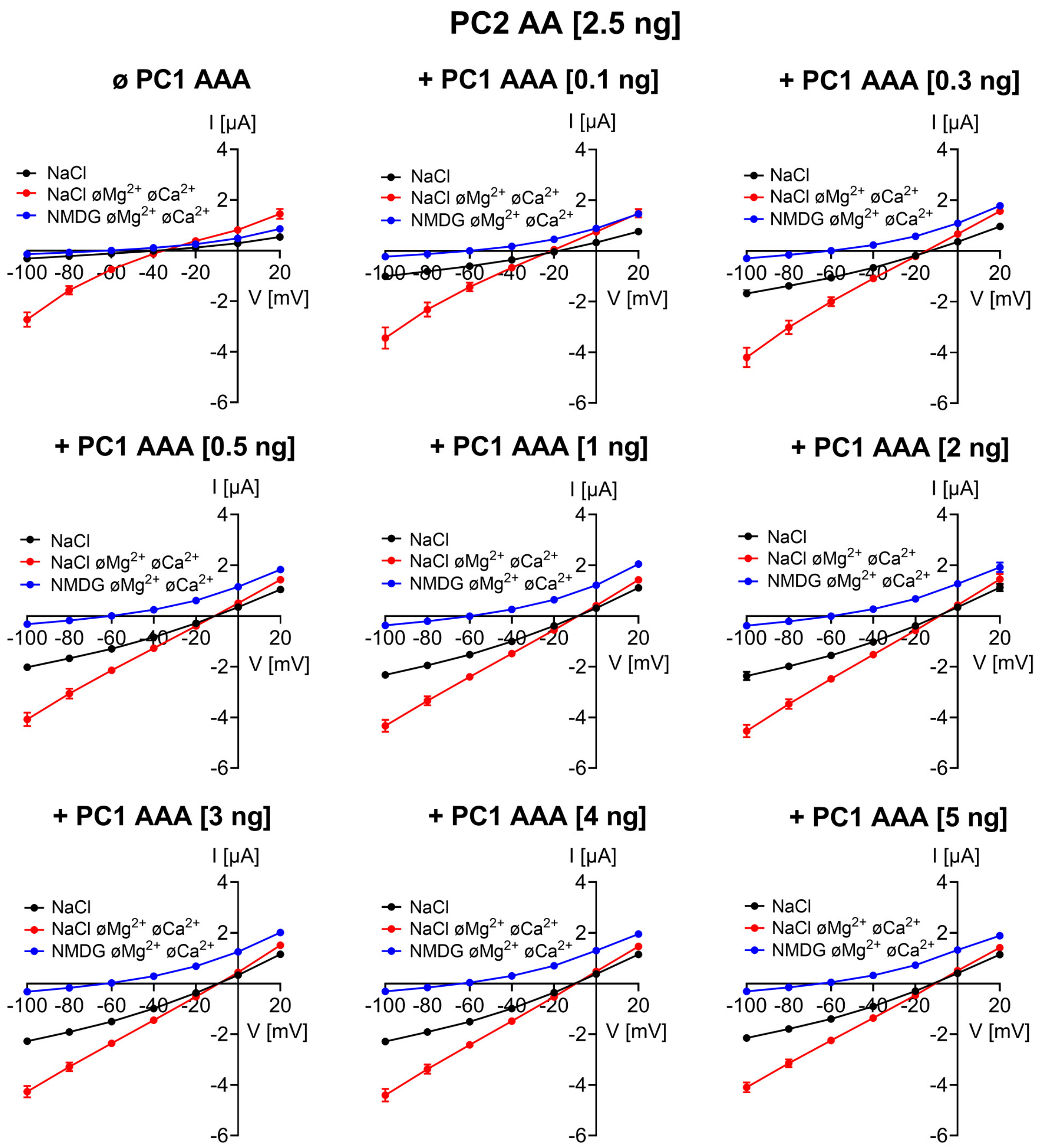


**Appendix Fig.S11 Average I/V plots from coexpression experiments using a fixed amount of PC2 AA and increasing amounts of PC1 AAA.**

Average I/V-plots (mean ± SEM; 13≤*n*≤27, N=2-3) obtained in different bath solutions using the same experimental approach as shown in Fig. 1 from oocytes co-injected with a constant amount of PC2 AA (2.5 ng) and a variable amount of PC1 AAA as indicated. Average I/V plots shown in Fig. 3 for NaCl bath solution without divalent cations (NaCl ø$Mg^{2+}$ø$Ca^{2+}$) are extracted from the I/V plots shown here.

| Metric | 6A70.pdb[a] | this study[b] |
|---|---|---|
| no. residues | 1786 | 1838 |
| no. atoms | 13237 | 15008 |
| MolProbability score[1] | 1.84 | 1.77 |
| CaBLAM outliers[2] | 4.75 % | 3.22 % |
| ADP (B-factors)[3] min/max/mean | 0.00/215.33/90.42 | 36.97/160.00/115.76 |
| $CC_{box}$[4] | 0.52 | 0.59 |
| $CC_{mask}$[5] | 0.59 | 0.74 |
| $CC_{volume}$[6] | 0.58 | 0.73 |
| $CC_{peaks}$[7] | 0.34 | 0.43 |

**Appendix Tab.S1 Comparison of the model-*versus*-data metrics for PC1/PC2 heterocomplex of the published (PDB ID: 6A70) and re-interpreted model (this study).**

For the comparison, the N-terminal domain (NTD, res. 3075-3657) from the original model was omitted in both models. Metrics were prepared using Python-based Hierarchical ENvironment for Integrated Xtallography (*Liebschner D. et al. Macromolecular structure determination using X-rays, neutrons and electrons: recent developments in Phenix. Acta Crystallogr D Struct Biol. 2019 75(Pt 10):861-877. doi: 10.1107/S2059798319011471*).

[a] Model residues of PC1: 3657-3752, 3783-4050, 4081-4120; PC2 (monomer 1): 219-291, 313-463, 471-694; PC2 (monomer 2): 219-293, 313-699; PC2 (monomer 3): 219-294, 305-699.
[b] Model residues of PC1: 3657-3749, 3783-3820, 3826-4122; PC2 (monomer 1): 216-294, 306-694; PC2 (monomer 2): 216-295, 304-694; PC2 (monomer 3): 216-295, 304-694.

[1] A log-weighted composite value of the clash-score, percentage Ramachandran not favored and percentage bad sidechain rotamers (lower = better).
[2] System to evaluate protein CA mainchain geometry in low-resolution structures.
[3] Atomic displacement parameter (ADP), uncertainty in atomic positions.
[4] Correlation coefficient, similarity of model and target maps.
[5] Correlation coefficient, fit of the atomic centers.
[6] Correlation coefficient, fit of the molecular envelope defined by the model map.
[7] Correlation coefficient, fit of the strongest peaks in the model and target maps.

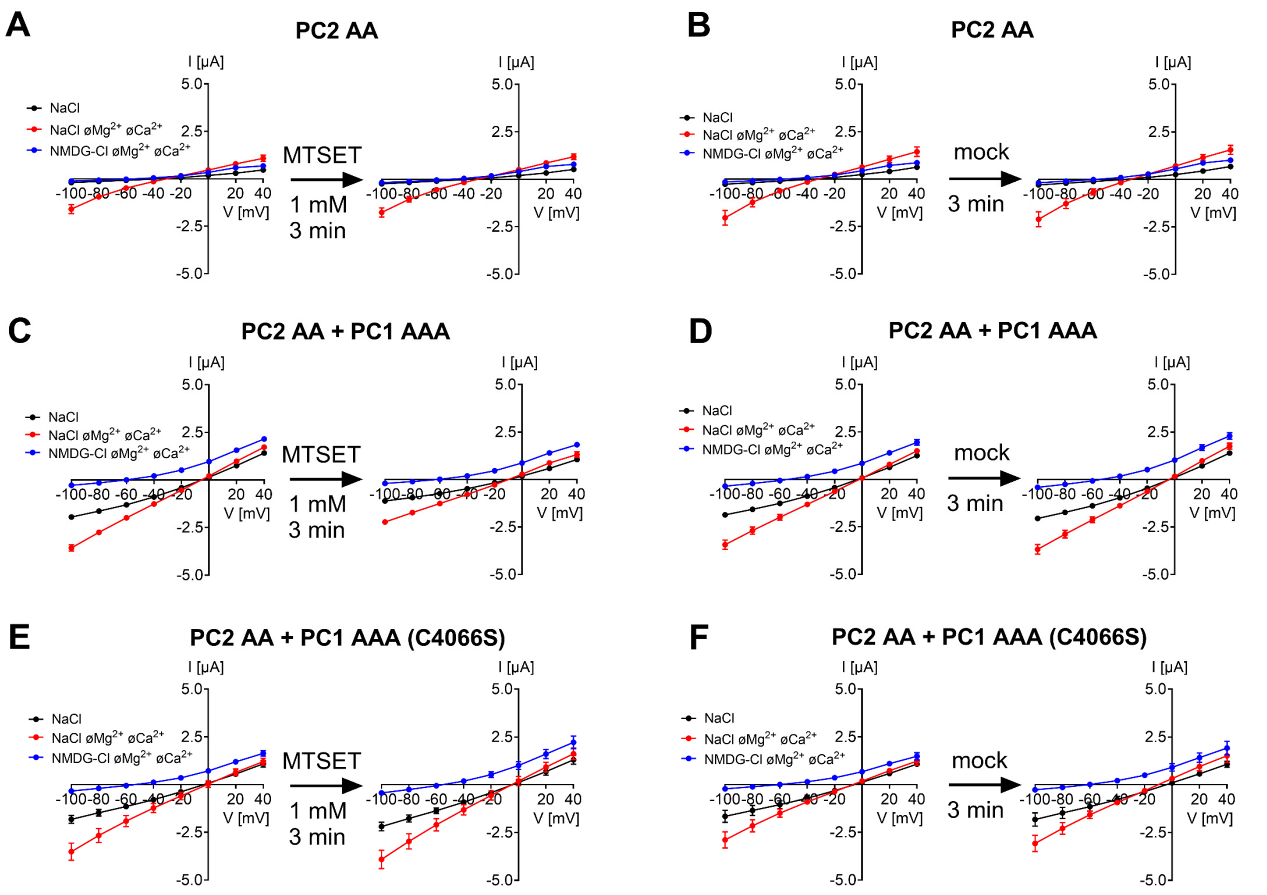


**Appendix Fig.S12 Sulfhydryl reagent MTSET probably inhibits PC2 AA + PC1 AAA heteromers through covalent modification of the pore-loop residue C4066**

Average I/V-plots (mean ± SEM; 8≤*n*≤12, N=2-3) obtained in different bath solutions using the same experimental approach as shown in Fig. 1 from oocytes expressing PC2 AA alone (***A***, ***B***), co-expressing PC2 AA and PC1 AAA (***C***, ***D***), or co-expressing PC2 AA and PC1 AAA with additional C4066S mutation (***E***, ***F***). In each individual oocyte currents were measured before and after 3 min incubation in NaCl bath solution supplemented with 1 mM MTSET (***A***, ***C***, ***E***). The oocyte was unclamped during the incubation time. Before the second current measurement, MTSET was washed out with NaCl bath solution. Impaling microelectrodes were not removed from the oocyte until the end of the experiment. Mock-treated control oocytes were incubated for 3 min in NaCl bath solution without MTSET (***B***, ***D***, ***F***). Average I/V plots shown in Fig. 4B for NaCl bath solution without divalent cations (NaCl øMg$^{2+}$øCa$^{2+}$) are extracted from the I/V plots shown here.
